# Ku suppresses RNA-mediated innate immune responses in human cells to accommodate primate-specific Alu expansion

**DOI:** 10.1101/2025.01.31.633084

**Authors:** Yimeng Zhu, Angelina Li, Suvrajit Maji, Brian J. Lee, Sophie M. Korn, Jake A. Gertie, Tyler J. Dorrity, Jianhua Wang, Kyle J. Wang, Amandine Pelletier, Daniel F. Moakley, Rachel D. Kelly, Antony B. Holmes, Raul Rabadan, David R. Edgell, Caroline Schild Poulter, Mauro Modesti, Anna-Lena Steckelberg, Eric A. Hendrickson, Hachung Chung, Chaolin Zhang, Shan Zha

## Abstract

Ku70 and Ku80 form Ku, a ring-shaped protein that initiates the non-homologous end-joining (NHEJ) DNA repair pathway.^1^ Specifically, Ku binds to double-stranded DNA (dsDNA) ends and recruits other NHEJ factors (*e.g.*, DNA-PKcs and LIG4). While Ku binds to double-stranded RNA (dsRNA)^2^ and traps mutated-DNA-PKcs on ribosomal RNA *in vivo,*^3,4^ the physiological significance of Ku-dsRNA interactions in otherwise wild-type cells remains elusive. Intriguingly, while dispensable for murine development,^5,6^ Ku is essential in human cells.^7^ Despite similar genome sizes, human cells express ∼100-fold more Ku than mouse cells, implying functions beyond NHEJ, possibly through a dose-sensitive interaction with dsRNA, which is ∼100 times weaker than with dsDNA.^2,8^ While investigating the essentiality of Ku in human cells, we found that depletion of Ku - unlike LIG4 - induces profound interferon (IFN) and NF-kB responses reliant on the dsRNA-sensor MDA5/RIG-I and adaptor MAVS. Prolonged Ku-degradation also activates other dsRNA-sensors, e.g. PKR that suppresses protein translation, and OAS/RNaseL that cleaves rRNAs and eventually induces growth arrest and cell death. MAVS, RIG-I, or MDA5 knockouts suppressed IFN signaling and, together with PKR knockouts, partially rescued Ku-depleted human cells. Ku-irCLIP analyses revealed that Ku binds to diverse dsRNA, predominantly stem-loops in primate-specific Alu elements^9^ at anti-sense orientation in introns and 3’-UTRs. Ku expression rose sharply in higher primates tightly correlating with Alu-expansion (r = 0.94/0.95). Together, our study identified a vital role of Ku in accommodating Alu-expansion in primates by mitigating a dsRNA-induced innate immune response, explaining the rise of Ku levels and its essentiality in human cells.

---

The NHEJ pathway repairs DNA double-strand breaks (DSBs) by directly ligating two DNA ends and plays a critical role in lymphocyte-specific gene rearrangements.^10^ NHEJ is initiated via the high-affinity binding of DSB ends by the Ku70-Ku80 heterodimeric ring (herein Ku), which then recruits other NHEJ factors, including Ligase 4 (LIG4) and the catalytic subunit of the DNA-dependent protein kinase (DNA-PKcs)^1^. While the role of Ku in NHEJ is well-characterized, accumulating evidence suggests that Ku has NHEJ-independent functions, especially in human cells. While dispensable for murine development,^5,6^ Ku, but not LIG4, is essential for the survival of all human cells tested.^7^ Ku70 and Ku80 (also called Ku86 in humans) depend on each other for protein stability. Analyses of the Dependency Map data (DepMap, 2022 Release 4)^11^ showed that both Ku80 (*XRCC5*) and Ku70 (*XRCC6*) are universally essential in >1000 human cancer cell lines tested via both CRISPR and RNAi **(Extended Data Fig. 1a-b).** Meanwhile, neither LIG4 nor DNA-PKcs are essential (**Extended Data Fig. 1c-d**). Second, despite having a similar genomic size, human cells examined have 50-100 fold more Ku **(Fig. 1a)** and DNA-PKcs^12^ protein than mouse cells. This is not due to limited antibody cross-activity, as XRCC5 and XRCC6 mRNA levels are 20–100 times higher in human compared to mouse cells **(Fig. 1b-c)**. The difference is even more striking when comparing the same cell type—naïve B cells **(Extended Data Fig. 1e-f)**. In contrast, LIG4 and its partner XRCC4 show comparable levels in human and mouse cells **(Fig. 1d and Extended Data Fig. 1g)**.

**Figure 1:**
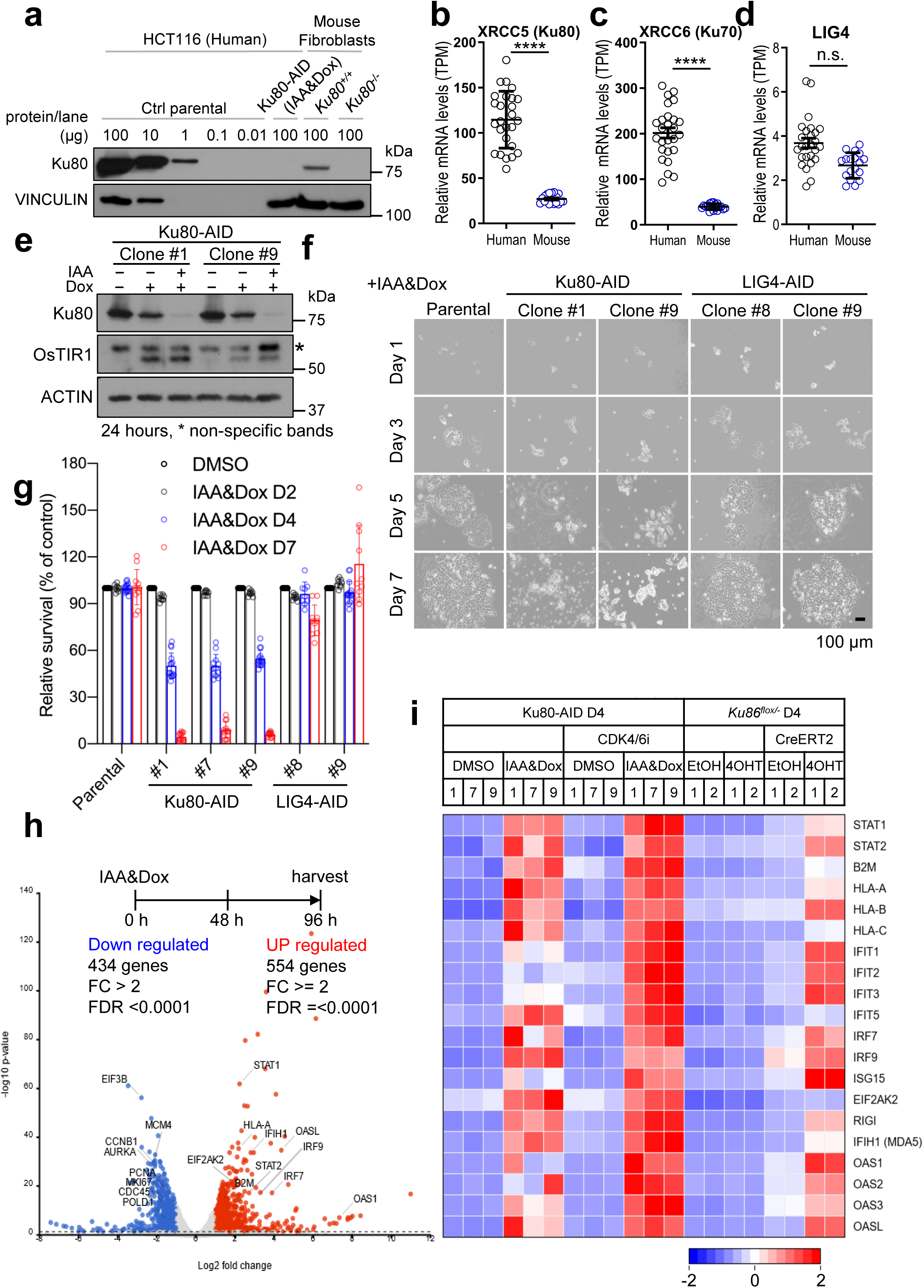
Ku expression is 100-fold higher in human cells, and degraded Ku leads to progressive lethality and activation of IFN. **a,** Western blot analyses of Ku80 protein in human colorectal cancer cell line HCT116 and immortalized murine embryonic fibroblasts (iMEFs). **b-d,** The normalized mRNA levels (transcript per million/TPM) for *XRCC5/Ku80* (b), *XRCC6/Ku70* (c), and *LIG4* (d) in human cell lines (HCT116, Hela) and primary mouse B cells, fibroblasts, and embryonic stem cells. The bars represent the mean and standard error of the mean (SEM) and unpaired student’s *t*-test *p*-value < 0.0001, ****. **e,** Doxycycline (Dox) and IAA (Auxin) successfully induced the degradation of AID-tagged Ku80 from two independent clones within 24 hr. *: a non-specific band. **f,** Representative images of parental, Ku80-AID, and LIG4-AID tagged HCT116 cells after Dox and IAA treatment. The pictures were taken at indicated times after treatment started. Representative images from each clone are shown. **g,** Relative viability (versus untreated) of parental cells, Ku80-AID, and LIG4-AID clones after IAA and Dox treatments measured by CellTiter-Glo luminescent cell viability assay. The plot represents the means and standard derivation from 3 independent experiments from each clone. The unpaired student’s *t*-test *p* < 0.0001 ****; *p* > 0.05 ns. **h,** Volcano plot of mRNA expression at 96 hr/4 day after induced Ku80 degradation. The diagram above shows the scheme of the experiment. Log2 fold change (FC) and negative Log10 *p*-value were plotted from DeSeq2 analyses of three independently derived Ku80-AID clones. The red and blue dots indicate significantly changed genes with FDR < 0.0001 and absolute fold change above 2. **i,** Heatmap of representative IFN-inducible genes. The color gradient reflects the Z-scores calculated from normalized TPM across the displayed samples: red represents higher expression levels relative to the mean, while blue indicates lower expression levels. Three independently derived Ku80-AID clones and two repeats of the *Ku86^flox/-^:CreERT2* and control *Ku86^flox/-^*clones are shown. For CDK4/6 inhibitor treatment, the inhibitors were added simultaneously with IAA and Dox, as Ku degradation requires 24 hr.

Ku forms a stable ring that distinguishes broken DNA ends from intact double-stranded DNA (dsDNA) by threading onto the DNA from its ends.^13^ Structural analyses showed that the Ku ring is sized to encircle 15-20 bp of dsDNA snugly, independent of sequence and end structures (*e.g*., blunt, small overhangs, and small stem-loops (SLs)),^13^ enabling a rapid and efficient response to a broad range of DSBs. In addition to DNA, Ku can also bind to dsRNA ends/SLs, albeit with ∼ 100-fold lower affinity.^8^ In *Saccharomyces cerevisiae*, yKu binds to an SL in the telomerase RNA template (TLC1) to facilitate telomere maintenance.^14,15^ Mammalian Ku also binds to the telomerase (TERT) RNA template^16,17^. However, cancer cell lines such as U2OS, CAL72, and SAOS2, which express nearly no TERT and maintain telomeres through the TERT-independent alternative lengthening of telomeres (ALT) mechanism, also depend on Ku for survival (**Extended Data Fig. 1h**), suggesting that Ku has another NHEJ-independent function in human cells beyond telomere maintenance. Notably, Ku and DNA-PKcs, but not other NHEJ factors (*e.g.*, LIG4, XRCC4), can be found in the RNA-rich nucleolus in a Pol-I transcription-dependent manner.^3^ Using mouse models, we previously reported that Ku-mediated recruitment and trapping of phosphorylation-defective or kinase-dead DNA-PKcs on the rRNAs and small nucleolar RNAs (snoRNA) compromise rRNA processing, protein translation, and eventually causes lethal anemia.^3^ However, the role of Ku and mutant DNA-PKcs in rRNA processing cannot explain the selective essentiality of Ku in human cells since ribosomal biogenesis is essential in all species. Ku deficiency also causes no anemia in mice^3^, leaving the physiological significance of Ku-RNA interaction in otherwise normal human cells unknown.

Given Ku’s relatively lower affinity for RNA, we sought to investigate the biological significance of Ku-RNA interactions in human cells, where Ku concentration is much higher. To circumvent the Ku-essentiality, we adapted the Auxin-Inducible Degron (AID) system^18^ and tagged both alleles of *XRCC5* (encoding Ku80) with AID in HCT116 cells (**Extended Data Fig. 2a-b**). We chose the HCT116 cell line because of its stable diploid karyotype, high targeting efficiency, *Tp53*-proficient, and telomerase-positive status.^19^ The doxycycline (Dox), which induces OsTIR1 expression, together with Auxin (IAA) efficiently degraded Ku80 by 24 hours (hr) **(Fig.1e)**. With the same strategy, we also generated several LIG4-AID clones as controls (**Extended Data Fig. 2c-d**). The attempt to degrade DNA-PKcs with the AID tag was unsuccessful, potentially due to the very large size of DNA-PKcs (4128 amino acids or ∼460 kDa). After IAA and Dox induction, Ku protein levels stayed low for at least nine days (**Extended Data Fig. 2e**). During this period, the cells progressively lost viability **(Fig. 1f-g)**, remaining largely unaffected on days 2 to 3, with minimal apoptosis observed by day 4 (**Extended Data Fig. 2f-g**), followed by a rapid decline from days 5 to 7, consistent with a positive feedback mechanism. In stark contrast to the loss of other essential genes involved in DNA repair and replication (*e.g.*, ATR, CtIP),^20,21^ Ku depletion only induced a partial G1/S cell-cycle arrest and mild replication fork stalling, as evidenced by BrdU-negative S-phase cells **(Extended Data Fig. 2h-i)**.

To understand the cause of viability loss, we collected RNA at Day 2 (48 hr), 4, and 6 after Ku-degradation. On Day 4 after Ku-degradation, before massive cell death, RNA-seq analyses from three independently derived Ku-AID clones identified ∼1000 genes with >2 folds absolute changes and FDR < 0.0001 **(Fig.1h)**. Consistent with growth arrest, the downregulated genes are frequently associated with proliferation (*e.g.*, MKi67, Cyclin B/CCNB1), DNA replication *(e.g*., MCM4, PCNA, POLD), and mitosis (*e.g.*, AURKA) (**Fig. 1h and Extended Data Table 1a**). Strikingly, the most upregulated pathways are interferon (IFN) signaling and, to a lesser extent, the related TNFα/NF-kB pathways **(Fig. 1h-i, and Extended Data Table 1b and Extended Data Fig. 3a and 3b)**. Moreover, several related inflammation-associated pathways (e.g., Inflammatory, IL-6-Jak-STAT3, IL-2 STAT5) are also significantly upregulated **(Extended Data Table 1b)**. IFNs are widely expressed cytokines with antiviral and growth-inhibitory effects, ^22^ and include three types - Type I (α, β), II (γ), and III (λ). While IFN-γ is produced primarily by active T cells and Nature Killer cells, most cell types can produce Type I IFNs and activate the NF-kB pathway in response to viral infection.^22^ Many IFN-inducible genes are activated by both Type I and Type II IFNs; however, a subset directly involved in limiting virus production, such as 2’,5’-oligoadenylate synthetases (OAS), and protein kinase R (PKR/EIF2AK2) are preferentially induced by Type I IFNs.^23^ Ku-degradation activates many IFN targets, including signaling mediators STAT1 and STAT2, transcriptional factors IFN-regulatory factor (IRF)7 and IRF9, viral RNA sensors (MDA5/IFIH1, RIG-I/DDX58), all components of the MHC I (HLAs and B2M), and notably Type I IFN selective targets OAS1-3 & OASL, and PKR implicated in growth arrest and cell death upon persistent virus infection (**Fig. 1i and Extended Data Table 2**). Other upregulated pathways also include apoptosis and checkpoint responses (**Fig. 1i-h and Extended Data Tables 1b and 2**). Given that growth defects at 4 day are likely indirect, we performed RNA-seq at 24 hr after Ku-degradation. While the IFN pathway is also significantly upregulated, albeit with more moderate fold changes (**Extended Data Fig. 3c and Extended Data Table 3**), the downregulated pathways at the early timepoint frequently involve RNA metabolism, from splicing to rRNA processing (**Extended Data Fig. 3d**). Given the implication of Ku^14,15^ and DNA-PKcs^24,25^ in DNA repair and telomere maintenance and the critical role of mitosis in promoting micronuclei formation and cGAS-STING mediated dsDNA-induced IFN and NF-kB activation,^26^ we tested whether proliferation is required for the IFN and NF-kB-induction upon Ku-degradation. CDK4/6 inhibitor (Palbociclib, CDKi) successfully arrested the cells in G1/G0, as measured by the diminished expression of replication-associated genes (**Extended Data Fig. 3e**). Yet, IFN signaling was still robustly induced upon Ku-degradation (**Fig. 1i, Extended Data Fig. 3f-h and Extended Data Table 4)** and followed by apoptosis on day 6 **(Extended Data Fig. 2f-g)**, suggesting that upon Ku degradation, both IFN and NF-kB activation and cell death can occur independent of mitosis or DNA replication. Mitosis seems to act as an accelerator instead. Once made, Type I IFNs bind to their receptors through autocrine and paracrine mechanisms to activate the Janus-activated kinase (JAK) family, which phosphorylates STAT1/2^22^. STAT1 and 2 are also transcriptional targets of the IFN cascade, forming a positive feedback loop^22^. In parallel, activation of NF-kB promotes the production of proinflammatory cytokines, which further amplify NF-kB signaling.^27^ Using quantitative RT-PCR, we validated the robust induction of IFN targets: ISG15, IFN-β, IFIT1, and STAT1, as well as the NF-kB targets: GADD45B, JUN, and PPP1R15A following Ku-degradation **(Fig.2a-d and Extended Data Fig. 3i-k)**. Western blotting further revealed robust STAT1(Y701) phosphorylation, cleavage of NF-kB p105 to p50, and the accumulation of IFN targets-STAT1, MDA5, and RIGI after Ku degradation (**Fig. 2e and Extended Data Fig. 3l)**.^22,26^ Consistent with the positive feedback of the IFN pathway, STAT1 phosphorylation first became evident on day 2 after Ku degradation and accumulated through days 4 to 9 **(Fig. 2f)**. Meanwhile, DMSO treatment alone, IAA/Dox treatment of the parental cell line, or degradation of LIG4, a core NHEJ factor, did not induce substantial STAT1 phosphorylation nor expression of IFN/NF-kB targets under the same conditions (**Fig. 2e and 2g)**, suggesting that Ku has a LIG4-independent and by extension, NHEJ-independent function, in suppressing innate immune responses.

**Figure 2:**
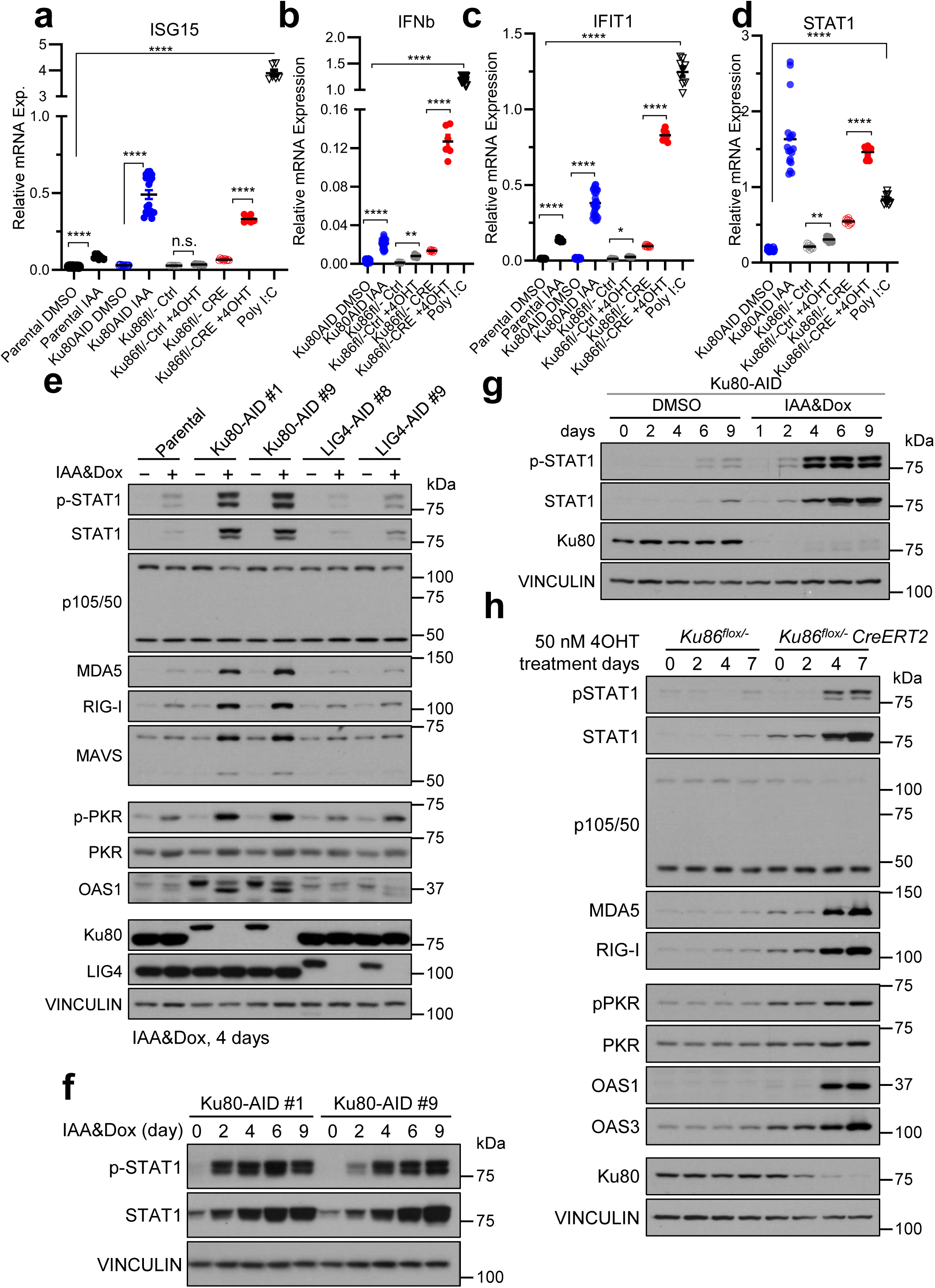
Ku degradation induces progressive and robust IFN signaling in multiple cell systems. **a-d,** Quantitative RT-PCR analyses of IFN-inducible genes (ISG15 (a), IFNβ (b), IFIT1(c), and STAT1(d)) were normalized to β-actin. The data include parental HCT116 with the AID system, three Ku80-AID clones (measured individually and combined), and replicates from *Ku86^flox/-^:CreERT2* and control *Ku86^flox/-^* clones. Bars indicate the mean ± SEM. ANOVA test significance:: *, p < 0.05; **, p < 0.01; ****, p < 0.0001; n.s., p > 0.05. **e,** Representative western blotting analyses of IFN and NF-kB pathway targets, and phosphorylation of PKR (T451) and STAT1 (Y701) at day 4 after inducing Ku80 or LIG4 degradation. **f and g,** Representative western blotting analyses of STAT1 and STAT1 phosphorylation at different days after Ku-degradation in multiple clones. DMSO treatment were included as control (g). Western blottings (e-g) were performed on at least three independent Ku-AID clones, with n ≥ 3 replicates. A representative result is shown. Quantification for NF-kB (p50/p100) ratio is at Extended Data Fig. 3l. **h,** Representative western blotting analyses of IFN and NF-kB pathways in *Ku86^flox/-^:CreERT2* and control *Ku86^flox/-^* clones treated with 4OHT (50nM). One out of four independent repeats were shown.

To ensure that the IFN/NF-kB activation is specifically caused by Ku degradation, and not by the IAA/Dox treatment itself, we acquired a set of HCT116 cells with genetically inducible Ku80 deletion. The *Ku86^flox/-^* HCT116 cells contain one deleted and one floxed Ku86 allele.^28^ An estrogen receptor (ER) fused Cre-recombinase (CreERT2) was introduced to generate the *Ku86^flox/-^:CreERT2* line, in which the ER ligand, 4-hydroxy tamoxifen (4OHT) triggers nuclear translocation of Cre and excision of the floxed Ku86 allele from the *Ku86^flox/-^:CreERT2*, but not control *Ku86^flox/-^* cells. We confirmed that the successful excision of the Ku86 flox allele resulted in a marked loss of viability **(Extended Data Fig. 4a-b)**. Since this genetic system removed the Ku86/XRCC5 gene (not protein), it takes longer (4-7 days rather than 24 hr in the AID system) to deplete Ku protein **(Fig. 2h)**. Nevertheless, RNA-seq analyses and qRT-PCR confirmed the robust upregulation of IFN and NF-kB pathways in 4OHT treated *Ku86^flox/-^:CreERT2*, but not control *Ku86^flox/-^* cells **(Fig. 1i**, **Fig. 2a-d and Extended Data Fig.4c-f, Extended Data Table 5).** Western blotting shows the phosphorylation of STAT1, cleavage of NF-kB p105, and the induction of STAT1, MDA5, RIG-I, and OASs at days 4 and 7 after Ku deletion **(Fig. 2h)**. Notably, this genetic system completely avoids doxycycline and IAA and has no measurable background IFN induction in the 4OHT treated control *Ku86^flox/-^*cells, supporting a Ku-dependent activation of IFN and NF-kB.

Since Ku is known to bind to both dsDNA and dsRNA, we asked which nucleic acid sensors activate the Type I IFN production^22^. While the Toll-like receptor pathway recognizes nucleic acids on the cell membrane and endosomes, cGAS senses cytosolic dsDNA and synthesizes cyclic GMP-AMP (cGAMP) that binds and activates STING^29^, and RIG-I and MDA5 detect dsRNA in the cytoplasm and activate MAVS on the mitochondrial membrane, to activate IFN and NF-kB signaling pathways^27,30^. Since Ku likely binds to self-dsDNA/dsRNA (i.e., non-viral) in this context, we asked whether cGAS-STING or MAVS is involved and found that HCT116 cells, like many other human cancer cell lines, do not express cGAS **(Extended Data Fig. 5a)** and cGAS negative human cancer cells still require Ku for survival **(Extended Data Fig. 5b)**. Meanwhile, CRISPR KO of MAVS in multiple clones completely abolished STAT1 phosphorylation and IFN activation validated by qRT-PCR (**Fig. 3a-b and Extended data Fig. 5c-h and Extended Data Table 2)**, suggesting that Ku suppresses dsRNA-mediated IFN and NF-kB activation. MAVS can be activated by MDA5, a sensor for long dsRNA, or by RIG-I that recognizes 22-200 bp dsRNA with a 5’-triphosphate or diphosphate group.^30^ RIG-I can also be activated by small self-RNA generated by RNaseL.^31^ CRISPR KO of RIG-I abolished most STAT1 phosphorylation with a partial, yet significant attenuation by MDA5 KO (**Fig. 3a and Extended Data Fig. 5h),** suggesting that Ku prevents the activation of multiple dsRNA sensors with a dominant role of RIG-I in HCT116 cells. We note that the expression of dsRNA sensors varies greatly among different human cell lines tested **(Extended Data Fig. 5i)**. Considering the sequence-independent nature of Ku-nucleic acid interaction and the contribution of both RIG-I and MDA5 in HCT116, different sensors might be dominant in different cell lines.

**Figure 3:**
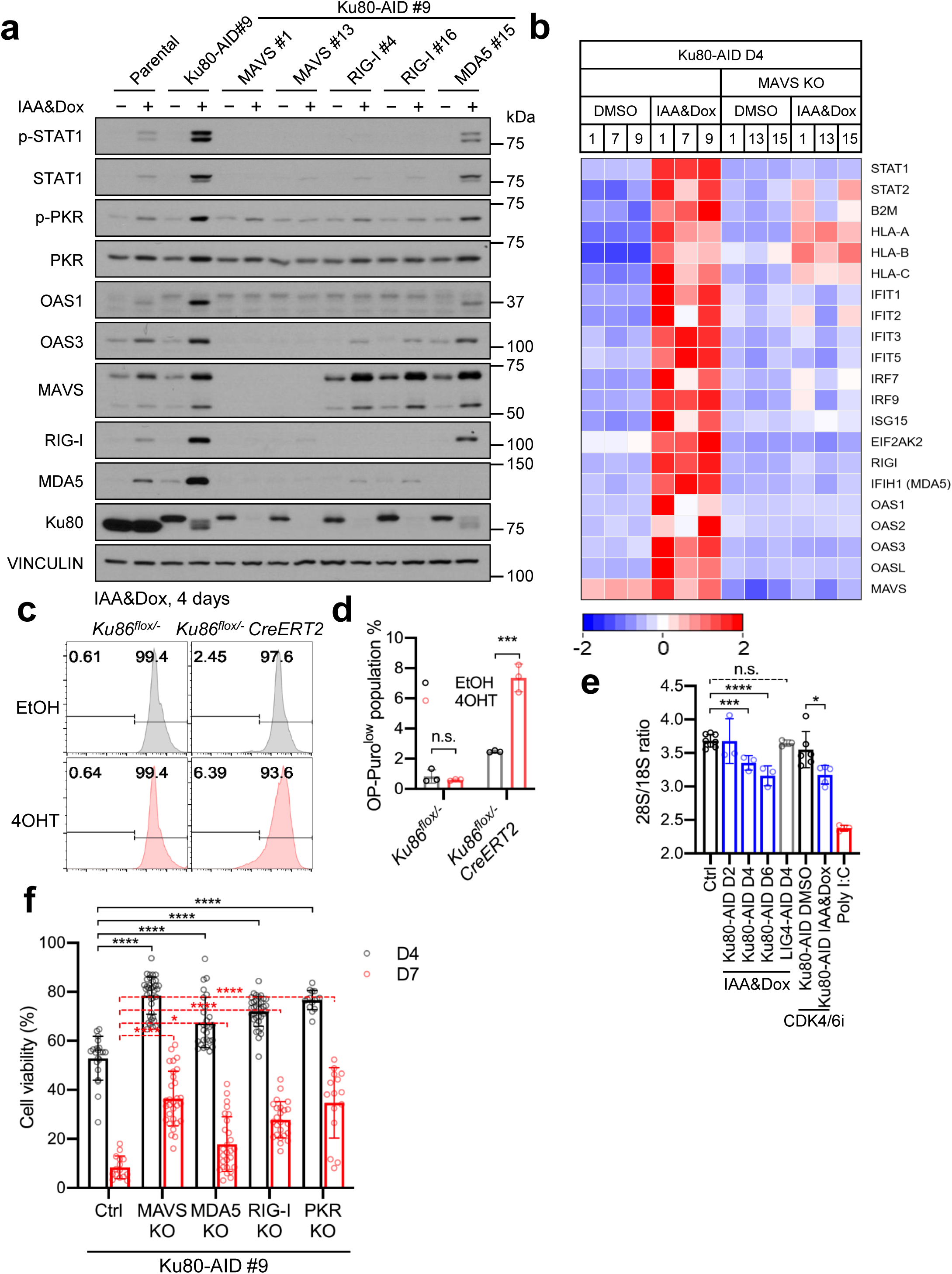
Ku degradation induces dsRNA- and MAVS-dependent IFN activation. **a** Representative western blotting analyses of IFN pathway markers in multiple independently derived MAVS, RIG-I, and MDA5 KO clones. Western blotting was performed on at least two, often three, independent clones for each KO, with n ≥ 3 replicates. A representative result is shown. Additional clones can be found in **Extended Data Fig. 5h. b,** Heatmap of representative IFN inducible genes. Three Ku80-AID clones and three independently derived MAVS KO clones are shown. The color gradient reflects the Z-scores calculated from normalized TPM across the displayed samples: red represents higher expression levels relative to the mean, while blue indicates lower expression levels. **c,** Representative OP-Puro analyses of *Ku86^flox/-^:CreERT2* and control *Ku86^flox/-^* clones after 4OHT treatment. **d,** Quantification of OP-Puro for protein translation changes after Ku-depletion. **e,** The 28S versus 18S rRNA ratio in DMSO treated (Ctrl) cells versus Ku or LIG4 degraded cells. At least three independent samples were measured at each time point. The average and the standard errors were plotted. The *p*-values were calculated via an unpaired student *t*-test, *p* < 0.05 *; *p* < 0.001***; *p* < 0.0001****; *p* > 0.05 ns. **f,** Loss of MDA5, MAVS, RIG-I or PKR all significantly rescues the viability of Ku-deficient cells. Several (n ≥ 3) independent CRISPR knockout clones were generated for each gene (except PKR with 2 clones), and the results were measured individually and plotted together. The *p*-values were calculated via a two-way ANOVA test, *p* < 0.0001 ****; *p* < 0.05 *.

In both parental and Ku80-AID HCT116 cells (without IAA/Dox), IFNβ up to 40 ng/ml did not cause massive cell death, while the dsRNA analog poly I:C at 10 and 100 ng/ml did **(Extended Data Fig. 6a-b)**. What triggers growth arrest and cell death? In cells deficient for ADAR1, a dsRNA-specific adenosine deaminase that converts adenosine to inosine, self dsRNA triggers growth arrest and cell death by activating PKR^32^ and OAS-RNaseL^33^. Upon binding to dsRNA (> 33 bp), PKR dimerizes, activates, and autophosphorylates itself and the translation initiation factor eIF2α.^34^ PKR phosphorylation inactivates eIF2α, shuts down protein synthesis, and compromises cellular viability.^34^ Given the role of PKR as an effector of dsRNA-induced cell death, we tested whether Ku-degradation activates PKR. Indeed, the loss of Ku, but not LIG4, induced robust PKR auto-phosphorylation by day 4 upon Ku-degradation or deletion **(Fig. 2e and 2h)**. Correspondingly, global protein synthesis, as measured by O-Propargyl-puromycin (OP-Puro) that is incorporated into nascent proteins^35^, also decreased in a significant population of Ku-depleted cells (**Fig. 3c-d, and Extended Data Fig. 6c)**. Like PKR, OAS family members are induced via IFN and activated by dsRNA.^33^ Active OAS enzymes join two ATPs to form the 2’-5’-linked adenosine, which acts as a 2^nd^ messenger to activate RNaseL, which in turn degrades viral and cellular RNAs, including rRNAs.^33,36^ Notably, RNaseL-cleaved self-RNAs can also activate RIG-I,^31^ forming a positive feedback loop. By day 4 after Ku-degradation or deletion, the mRNA and the protein levels of multiple OAS family members (OAS1-3 and OASL) increased significantly in both cycling and G1/G0 arreseted cells treated with CDK4/6 inhibitor **(Fig. 1i, 2e, 2h and Extended Data Fig. 6d)**.

Correspondingly, degradation of rRNA, especially the largest 28S rRNA, became apparent at 4 days and progressed at 6 days after Ku-depletion in both cycling and arrested cells and upon Ku-deletion **(Fig. 3e and Extended Data Fig. 6e-g).** Importantly, LIG4 degradation did not cause any measurable rRNA degradation **(Fig. 3e)**.

To test whether Ku induces innate immune responses in different human cells, we analyzed recently published Ku70 knockout (KO) HEK293 lines sustained by Dox-inducible expression of ectopic Ku70.^37^ In these HEK293 clones, Dox led to approximately 10-fold overexpression of Ku70 mRNA **(Extended Data Fig. 7a)**. While the Ku70 protein level remains normal due to the requirement for endogenous Ku80 protein for its stability, it took 6–8 days after Dox withdrawal for the excess Ku70 mRNA to be diluted and for Ku70 protein levels to be depleted **(Extended Data Fig. 7b)**. Correspondingly, a relatively moderate loss of viability was noted at day 7 (**Extended Data Fig. 7c**, ∼60% versus ∼5% in AID or the flox systems). Nevertheless, RNA-seq analyses on Day 8 after Dox withdrawal from all three clones (Sal1, SB and T1) revealed an inflammatory signature dominated by the NF-kB with moderate significance for IFN signaling **(Extended Data Fig. 7d-e, Extended data Table 6)**, which was validated by qRT-PCR and Western blotting **(Extended Data Fig. 3i-k and 7b)**. While it is unclear why NF-kB is dominant in HEK293 cells, MAVS activates both IFN and NF-kB.^27^ Even after IFNβ induction, HEK293 cells cannot express IRF7, critical for the type I IFN production and amplification of the IFN signaling **(Extended Data Fig. 7f and 5i)**. Importantly, despite the lack of prominent IFN activation, Ku depletion in HEK293 cells induced marked phosphorylation of PKR, reduced protein translation **(Extended Data Fig. 7b and 7g)** and induced significant degradation of 28S rRNA at day 8 **(Extended Data Fig. 7h)**, similar to HCT116 cells.

Given that OASs and PKR as well as RIG-I and MDA5 are transcriptional targets of IFN and IFN signaling is amplified through positive feedback after Ku degradation **(Fig. 1i, 2e, and 2h)**, we tested whether deletion of MDA5, RIGI, or MAVS individually can rescue the Ku-deficient human cells. Several MDA5, RIG-I, MAVS or PKR knockout clones were generated from Ku-AID-tagged cells **(Fig. 3a and Extended Data Fig. 5h and 8a)**. Deletion of MAVS and RIG-I, and to a lesser extent MDA5, blunted IFN activation after Ku-degradation (**Fig. 3a and Extended Data Fig. 5h)**. Deletion of PKR did not affect STAT1 activation, consistent with its role as a downstream effector **(Extended Data Fig. 8a)**. Nevertheless, MDA5, MAVS, RIG-I or PKR KO all significantly, albeit partially, rescued the viability of Ku-degraded cells (**Fig. 3f)**. The multipronged nature of dsRNA-induced cell death following Ku degradation aligns with Ku’s sequence-independent nucleic acid binding and its suppression of diverse dsRNA sensors, highlighting Ku’s essential role in all human cells examined. These findings suggest that Ku supports human cell viability through a LIG4- and NHEJ-independent mechanism by suppressing dsRNA-induced innate immune responses, including the activation of PKR and OAS/RNaseL, and potentially interfering with other RNA metabolic processes.

What are the human-specific dsRNAs that activate IFN upon Ku degradation? To determine the full spectrum of cellular RNAs bound by Ku in human cells, we reanalyzed our previously generated infrared crosslinking and immunoprecipitation (irCLIP) data.^3,38^ While our previous analyses focused on RNAs bounded by both Ku and DNA-PKcs within 30 nt from each other and identified rRNA and U3 snoRNA,^3^ here we analyzed RNAs bounded by Ku regardless of DNA-PKcs. Since the binding patterns of the two biological replicates are consistent with each other, we pooled the two replicates together for the analyses. More than 60% of Ku-irCLIP tags mapped to introns of mRNA/ncRNAs (**Fig. 4a and Extended Data Table 7**), while only ∼20% mapped to exonic sequences or extended 5’-/3’-UTR regions, consistent with widespread Ku-RNA interactions in the nucleus. Further analyses suggested that nearly 70% of Ku irCLIP tags reside in repetitive elements, with 56% of those tags in the short interspersed nuclear elements (SINEs), followed by 12% in the long interspersed nuclear elements (LINEs) and 11% in the rRNA (**Fig. 4b**). This striking enrichment in repetitive elements, especially SINEs, is not typically seen for canonical nuclear RNA-binding proteins (RBPs) (*e.g*., PTBP1 and RBFOX2**, Extended Data Fig. 9a-b)**, but shared by DNA-PKcs **(Extended Data Fig. 9c-d).** SINEs are nonautonomous retrotransposons that do not encode a functional reverse transcriptase.^39^ While LINEs predominantly reside in heterochromatin regions, SINEs are often found within or near gene-rich areas, favoring highly expressed housekeeping genes over tissue-specific genes.^40^ Thus, SINEs can influence multiple aspects of gene regulation, including transcription and RNA splicing. Approximately 13% of the human genome encodes SINEs,^39^ including ∼0.6 million copies of mammalian-wide interspersed repeat (MIR) elements and ∼1.2 million copies of Alu elements, named after the Alu I endonuclease.^39^ While the mouse genome contains limited MIR, B1, and B2 SINEs, Alu elements are primate-specific and underwent significant expansion between the lemur and marmoset by introduction of the AluS sub-families.^9^ AluS shares >85% sequence identity with the most ancient AluJ family and ∼90% with the youngest AluY family.^9^ The frequency of Ku- or DNA-PKcs-bound Alu elements of different subfamilies generally correlates with their abundance in the genome, showing no strong bias for specific Alu subfamilies. One exception is that the youngest AluY subfamily that is found less than expected in Ku and DNA-PKcs irCLIP targets **(Extended Data Fig. 9e-f)**. Consistent with the primate-specific expansion of Alu, we found that Ku protein levels are not only high in human cells but also in other non-human primates (*e.g.,* Cos7 derived from African green monkey/*Cercopithecus aethiops*), while low in non-primate mammals (*e.g.*, dogs, horses) (**Extended Data Fig. 8b).** Re-analyses of published RNA-seq data showed that Ku mRNA expression is ∼10-100 fold lower in non-primates, and prosimian primates (*e.g.*, lemur^41^)-before the major Alu expansion occurred -as compared to monkey (*e.g.*, rhesus macaque) and human (**Extended Data Fig. 8c-d)**. On the other hand, ADAR1 expression is only 3-fold higher in humans than in mice (**Extended Data Fig. 8e)**. Comprehensive analyses within primates showed that the expression of XRCC5 (encoding Ku80) and XRCC6 (encoding Ku70) tightly correlate with Alu copy number expansion (Pearson correlation r = 0.94 and 0.95, respectively). In contrast, no significant correlation was observed for LIG4 (R = −0.18), and LIG4 levels remained consistently low across all 10 primate species examined (**Fig. 4c and Extended Data Table 8**).

**Figure 4:**
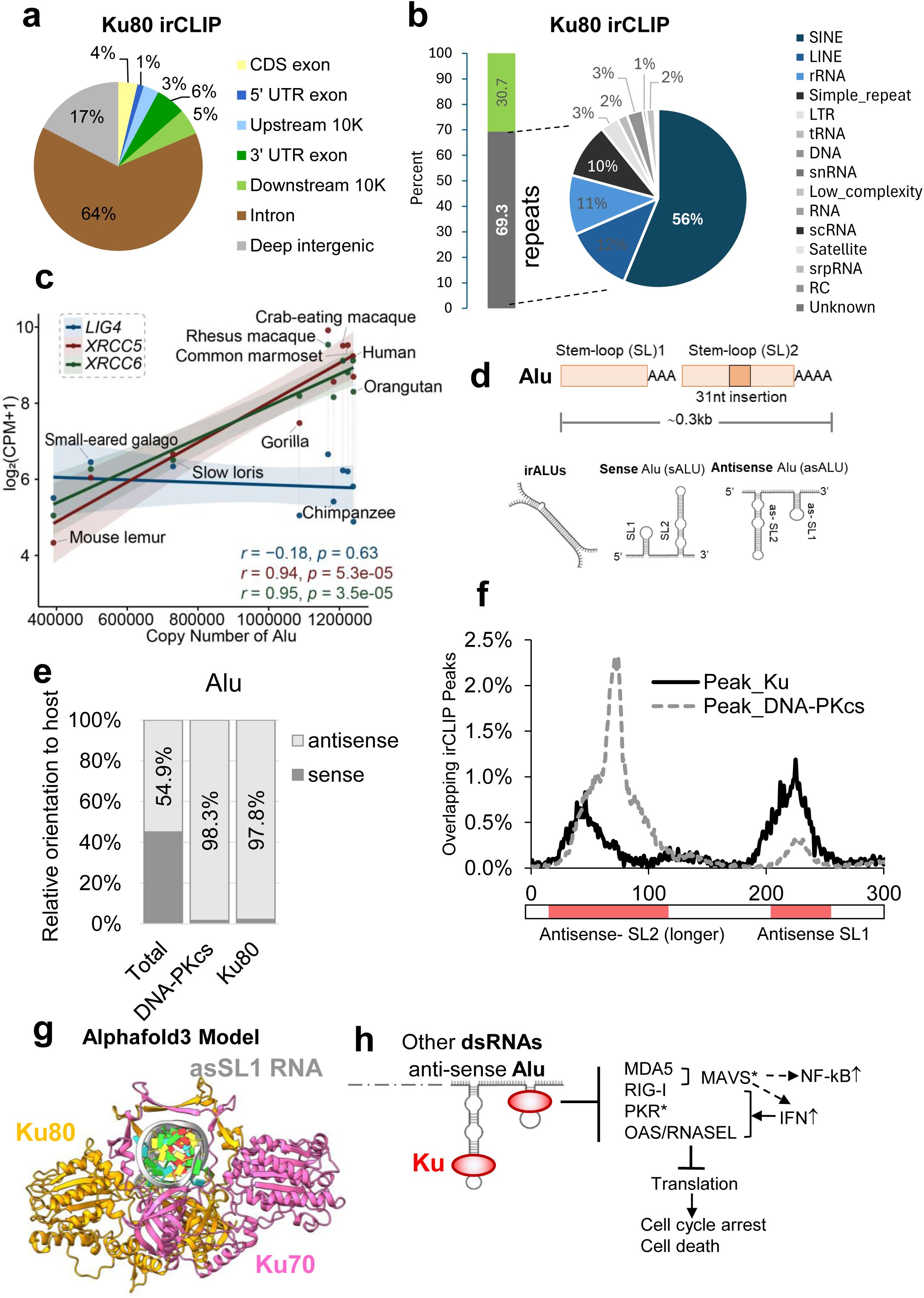
Ku binds to monomeric Alu in antisense stem-loops in introns and 3’-UTRs. **a,** Distribution of Ku-irCLIP tags. The results from two biological repeats were pooled together. **b,** Distribution of Ku-irCLIP tags among different types of repetitive elements. **c,** Evolution correlation of XRCC5 (red), XRCC6 (green), and LIG4 (blue) expression versus the Alu copy number expansion among primates. The correlations were assessed using Pearson’s correlation coefficient. **d,** Diagram of Alu elements and the folded RNA-secondary structure formed by the invert repeated Alus (irAlus), the sense and antisense monomeric Alu. **e,** Sense and antisense distribution of Alu in the genome versus those bounded by Ku or DNA-PKcs in irCLIP. **f,** The distribution of irCLIP peaks at each position in antisense Alu. The red boxes at the bottom indicate the antisense SL2 (longer) and SL1 (shorter). **g,** The Alphafold3 structural model of human Ku70 (pink) and Ku80 (yellow) bound to asSL1 of Alu RNA. All 5 top predictions show very similar configurations and the top predictions are shown here. **h,** A working model shows that Ku-Alu binding might prevent irAlu annealing, limit nuclear export of dsRNA, and directly compete with PKR and OASes to suppress IFN-activation and PKR and OASes-mediated cell growth arrest and cell death.

Alu elements are ∼300 nt long and contain the head-to-tail fusion of two distinct 7SL RNA monomers, a component of the signal recognition particle.^9^ Thus, each Alu contains two stem-loops – SL1 and SL2 – flanking an A-rich linker. A 31 nt insertion in the middle of SL2 makes SL2 longer than SL1 (**Fig. 4d**). Alu can form immunogenic dsRNA through at least two mechanisms. Short dsRNAs (22-40 bp) can form in monomeric Alus in both sense and antisense (as) strands depending on the relative orientation of the Alu and its host gene transcript (**Fig. 4d**). Alternatively, nearby inverted repeat Alus (irAlu) can pair with each other and form long dsRNA (**Fig. 4d**).^42^ The irAlu RNAs have been proposed to trigger pathological MDA5 activation in Aicardi-Goutières Syndrome (AGS), a type I interferonopathy^42,43^. Alu elements are also the primary targets of ADAR-mediated A-to-I editing in human cells, which introduces mismatches to suppress long dsRNA formation.^32^ ADAR deficiency also causes AGS.^43^

To understand how Ku suppresses Alu-induced type I interferonopathies, we analyzed the Ku irCLIP binding sites within Alus. **First**, to our surprise, neither Ku nor DNA-PKcs showed significant enrichment at the previously characterized immunogenic irAlus^42^; instead, irAlu is relatively depleted among the Ku or DNA-PKcs irCLIP targets as compared to the non-irAlus (Ku: 0.9% vs 2.1%, *p* = 2.077e-06; DNA-PKcs: 3.8% versus 7.1%, *p* = 1.4e-12; Fisher’s exact test; **Extended Data Fig. 10a-b**), prompting us to analyze Ku binding in monomeric Alu elements independent of the presence of irAlu pairs. **Second**, while Alu elements in the host gene can be transcribed in either sense or antisense orientations (45% sense versus 55% antisense, **Fig. 4d-e**), strikingly ∼98% of irCLIP target sites in Alu for Ku or DNA-PKcs are in the antisense orientation (asAlu) (**Fig. 4e and Extended Data Table 9**). Moreover, Ku and DNA-PKcs binding shows no major orientation bias at LINE elements **(Extended Data Fig. 10c)**. The overwhelming preference of Ku for asAlu suggests that it primarily interacts with monomeric Alu elements. Ku binding to the monomeric Alu SLs likely stabilizes the SL folding within individual Alu, thereby preventing adjacent Alus, including irAlus, from annealing into longer dsRNAs. Alternatively, due to Ku’s ring-shaped structure and its requirement to thread onto dsRNA from the ends, long irAlus may be less favored for Ku binding. **Third,** given the high sequence/structural conservation among Alu subfamilies and the lack of strong Alu-subfamily bias for Ku and DNA-PKcs binding **(Extended Data Fig. 9e-f),** we overlaid the irCLIP peaks on all asAlus to identify Ku and DNA-PKcs binding sites within monomeric asAlu elements. The results revealed that Ku and DNA-PKcs preferentially bind at the two predicted SLs in asAlu (**Fig. 4f and Extended Data Fig. 10d-e**), supporting the monomer binding mode and consistent with Ku’s known preference for dsRNAs, including SLs. Furthermore, the protein-RNA crosslink sites determined via crosslinking-induced mutation site (CIMS) and truncation site (CITS) analyses^44,45^ are also enriched at the two SLs upstream of the peak positions, which is consistent with the known preference for UV-crosslinking at nucleotides in bulges or proximal single-stranded regions **(Extended Data Fig. 11 and Extended Data Tables 10-11)**. The analyses also revealed a preference for DNA-PKcs binding at the longer asSL2 (30-35 bp) over the shorter asSL1 (20-25 bp) **(Extended Data Fig. 10e and 11)**. As an example, the transcript of MDM4, a negative regulator of Tp53, contains several asAlu elements in the 3’-UTR^46^. Both Ku and DNA-PKcs irCLIP sites were identified with a preference of Ku at asSL1 and DNA-PKcs at the predicted longer asSL2 **(Extended Data Fig. 12)**. **Extended Data Fig. 13** shows another example *LOC284454* with a similar pattern. Notably, two nearby microRNAs, Mir23A and Mir27A, are also bound by Ku and DNA-PKcs, but the pattern is similar for Ku versus DNA-PKcs. This suggests that the antisense orientation bias and the preference for asSL1 over asSL2 observed for Ku may be unique for Alu elements.

To validate Ku binding to Alu SLs, we measured the affinities of purified Ku to Alu SL1, asSL1, and a control perfectly paired dsDNA SL using fluorescence anisotropy. The results confirmed that Ku interacts directly with isolated Alu SL1 and asSL1 with an affinity that is approximately 10x lower than binding to the control dsDNA **(Extended Data Fig. 14)**. Ku showed similar affinities for SL1 and asSL1, consistent with sequence-independent interaction, suggesting that additional factors beyond the SL sequence tested here (*e.g.*, other RNA binding partners and flanking RNA sequences) dictate the antisense preference *in vivo*. Structural studies showed that the Ku encircles perfectly paired B-form dsDNA with a footprint spanning 15-20 bp^13,47^ **(Extended Data Fig. 8f-g)**. Alphafold3 modeling indicates that the core of human Ku encircles the asSL1 of Alu RNA in a similar configuration **(Fig. 4g and Extended Data Fig. 8f-g)**. A-form RNA has narrower and deeper grooves and is more compact (11 bp/helical turn) than B-form DNA (10 bp/helical turn). The modeling suggests that asSL1 slightly melts about a bulge (U19 and U21) to accommodate Ku binding, similar to what has been observed in the structure of yKu bound to TLC1 SL^14^ **(Extended Data Fig. 8f and 8h)**. On dsDNA, the addition of DNA-PKcs pushes Ku to rotate inward ∼10 bp^47^. Together, Ku and DNA-PKcs occupy ∼30 bp, which might explain the preference of DNA-PKcs for the longer asSL2 (∼30 bp). Alternatively, DNA-PKcs might bind to asSL2 independent of Ku since the A-form dsRNA might be challenging for the rotation of the Ku ring that snuggly fits the B-form dsDNA. Consistent with the latter model, we saw little correlation between Ku and DNA-PKcs irCLIP at the antisense Alus **(Extended Data Fig. 15 and Extended Data Table 10)**.

What is the impact of Ku binding on the Alu-containing mRNA? Ku-degradation did not consistently affect the levels of mRNA with experimentally defined Ku-irCLIP sites **(Extended Data Fig. 10f and Extended Data Table 12)** or those with genome-annotated Alus in their introns and 3’-UTR **(Extended Data Fig. 10g and Table 13)**. This suggests that Ku may bind to excised introns, which make up the majority of nuclear RNA and frequently contain Alu elements. Using the J2 antibody that recognizes dsRNA (> 40nt)^48^, we also did not observe major changes in dsRNA levels after Ku-degradation **(Extended Data Fig. 16a-b)**. While J2 staining might not be sensitive enough and has a size requirement (> 40nt), it does give a much higher signal in human HCT116 cells than murine embryonic fibroblasts (MEFs) and was completely eliminated by dsRNase III treatment, but not DNase or single strand RNase **(Extended Data Fig. 16c-e)**. Finally, we noted that with the 100-fold increase of Ku protein concentration in human cells, Ku does not only bind to asAlu RNA, but also to many other structured RNAs, including microRNA and tRNA **(Fig. 4b and Extended Data Fig. 13 and 17)**. Similar to its binding to Alu, Ku preferentially binds the SLs on tRNA with crosslink sites detected mostly frequently in the T͸C loop and the anti-codon loop **(Extended Data Fig. 18a-e and Extended Data Table 7)**. The diversity of RNAs bound by Ku in human cells corresponds with the activation of divergent RNA sensors with distinct sequence and structure preferences upon Ku depletion. Together, these data support a model in which Ku-binding shields or sequesters diverse nuclear dsRNAs, especially primate-specific Alu SLs at introns and 3’-UTRs, from various dsRNA sensors including RIG-I, MDA5, and, perhaps more importantly, PKR and OAS family members to accommodate Alu expansion in higher primates.

The essentiality of Ku in human cells and its high concentration have long been puzzling. Using engineered human cells with inducible rapid Ku degradation, we demonstrate that Ku loss triggers progressive cell death, with a slow onset followed by rapid collapse over ∼8 days, independent of cell division. This is distinct from telomere crisis (> 100 doublings) or replication-associated death within 2-3 divisions (e.g., ATR or CtIP loss). These kinetics resemble that of an uncontrolled viral infection and are consistent with positive feedback, characteristic of IFN signaling. Mechanistically, we uncovered an unexpected primate-specific role of Ku in accommodating Alu expansion by suppressing dsRNA-induced IFN and NF-κB activation and preventing PKR- and OAS-mediated growth arrest and cell death **(Fig. 4h)**. Specifically, we propose that the high Ku expression in human cells, complementing its relatively low affinity to dsRNA, allows Ku to bind to diverse dsRNA species, including the conserved asSLs in Alu elements at introns and 3’-UTRs, as well as tRNA and other structured RNAs. Ku binding suppresses IFN, NF-kB activation and prevents immunogenic RNA-induced cell death through potentially three mechanisms: 1) preventing the annealing of long dsRNA, including irAlu that activates MDA5, by reenforcing short SLs within individual Alu via Ku ring, 2) directly competing with RIG-I, OAS and PKR on RNA fragments, including shorter Alu asSLs, to prevent their activation, and 3) sequestering dsRNA in the nucleus away from the cytoplasm, where dsRNA sensors are abundant and primed for activation **(Fig. 4h)**. While MDA5 and MAVS primarily reside in the cytoplasm, a fraction of PKR and OASes and the abundant ADAR1-110p localize in the nucleus and can directly compete with Ku for dsRNA. While Ku is primarily a nuclear protein, when the nuclear membrane breaks down transiently during mitosis, Ku could limit the diffusion of free nuclear RNAs, explaining why mitosis is not essential but can accelerate cell death after Ku-depletion (Extended Data Fig. 2g). While HCT116 cells robustly activate the IFN and NF-κB pathways upon Ku depletion, leading to increased expression of dsRNA sensors such as PKR and OAS that accelerate cell death, autocrine IFN signaling is likely neither required nor sufficient for this process. Instead, the direct activation of PKR, OAS, and potentially other sensors is the primary cause of cell death.While we emphasize Ku’s novel RNA-dependent role and conducted our study in cGAS negative HCT116 and HEK293 cells^29^, complete Ku deficiency in human cells also compromises NHEJ, which could fuel IFN and NF-kB activation via the cGAS-STING pathway, exacerbating the growth arrest and cell death. Due to the high levels of Ku in human cells, complete or near-complete depletion of Ku may be required to significantly impair NHEJ.

We have focused on Alu, the most abundant primate-specific dsRNA, whose expansion in primates is closely associated with increased Ku expression, to elucidate Ku’s essential role in human cells. Compared to LINEs, Alu elements are more commonly found within gene bodies and actively transcribed regions, which explains their high abundance in RNA, particularly in introns and 3’-UTRs. However, the ring shape of Ku and its sequence-independent binding to dsRNA enable it to interact with a broad range of endogenous dsRNAs, including rRNA intermediates^3^, tRNA, and microRNAs **(Fig. 4b and Extended Data Fig. 13, 17 and 18)**. The role of Ku in suppressing multiple dsRNA sensors likely explains why the knockout of any single sensor can only partially restore viability and why Ku is essential in over 1000 human cell cancer lines tested. What is the impact of Ku binding on RNA? While our data suggests that Ku does not have a global impact on mRNA expression and dsRNA content, Alu has been implicated in various RNA processing events, including, but not limited to, the nuclear retention of long non-coding RNAs (lncRNAs),^49^ splicing,^50^ microRNA binding, and translation control. Whether Ku contributes to a subset of these Alu-mediated functions remains elusive. While this work established a critical role of Ku in coping with Alu expansion by suppressing dsRNA-mediated innate immune responses, future investigation is needed to establish 1) the functional interaction of Ku with other RNA binding proteins and RNA processing pathways, 2) how RNA-dependent assembly of Ku and DNA-PKcs differs from those on DNA, 3) what the role of DNA-PKcs on Alu expansion and rise of Ku levels are, and 4) how Ku assembly collaborates with other RNA modifiers, such as ADAR1 to ensure Alu-mediated cellular function and Alu tolerance while maintaining a sensitive anti-viral defense system.

## Acknowledgments

We thank Dr. Masato Kanemaki for providing the HCT116 OsTIR1 cell line, Dr. Katheryn Meeks for providing cell lines derived from dogs and horses, and Dr. Riccardo Dalla-Favera for providing transcriptional data from human primary B cells. We thank Drs. Sun Hur and Cheng-Zhong Zhang from Harvard Medical School for sharing the irAlu and non-irAlu annotations. We thank Drs. Wei Yang at NIH, Xuebing Wu, Richard Baer, and Katia Basso at Columbia University for their helpful discussions, Dr. Teresa Palomero for information on the JAK signaling pathway, and Dr. Alberto Ciccia for cGAS-STING/IFN pathway detection reagents. We thank Drs. Sara Prezetocka and Jan Karlseder for helpful discussion and advice about ZBP1, and thank Dr. Stephen P. Goff for advice about virus RNA SLs during the revision. Due to space constraints, we often cited reviews rather than original publications. We apologize to the colleagues whose original works were not cited here.

## Author contributions

The Zha group uncovered the role of Ku in suppressing the RNA-mediated innate immune response in human cells and generated the Ku-irCLIP data. The Zhang laboratory conducted the irCLIP data analyses and uncovered the preferential binding of Ku to Alu elements. Specifically, Y.Z. and S.Z. conceived the initial idea to characterize the essentiality of Ku in human cells and designed KU degron experiments. Y.Z. generated the Ku80-AID cells, and A.L helped Y.Z generate LIG4-AID control cells and carried out the qPCR analyses with help from B.L.. Y.Z. and S.Z. designed, and Y.Z carried out all the cellular experiments, western blot analyses and collected all the RNA for the RNA-seq analyses with help from A.L. and B.L. B.L, S.Z., and A.L. analyzed the RNA-seq data with help from A.H. B.L. carried out the DeSeq2 analyses and generated all the final heatmaps, volcano plots, and pathway analyses related RNA-seq. A.H provided the human B cell transcriptome data and developed the software for generating heatmap and pathway analyses used in the paper. S.M. and C.Z analyzed the irCLIP data. DFM explored the alternative mechanisms of RNA mediated Ku-function. A.M. and M.M. purified the human Ku proteins used in the binding assay. A.S., S.M.K, and Y.Z. designed and S.M.K. and Y.Z. performed the fluorescence anisotropy experiments. K.W. performed the structural modeling of Ku with dsRNA and dsDNA and generated the related figures. J.W. in RR’s lab carried out the evoluation analyses of Ku expression and Alu expansion and generated the related figures. E.A.H. provided the *Ku86^flox/-^:CreERT2* and control *Ku86^flox/-^* HCT116 cell lines and discussed impact of Ku-depletion in human cells. R.D.K, D.R.E. and C.SP. provided the Ku70 KO HEK293 cell lines with Dox-inducible Ku70 expression. H.C. designed the J2 antibody experiments, and J.A.G. and T.D. performed and analyzed the J2 antibody staining. H.C. provided significant insights and knowledge on experimental design and the characterization of RNA-dependent innate immune responses and helped us interpret the data. S.Z. prepared the initial draft with help from Y.Z. and C.Z., and all authors contributed to the writing of the final manuscript.

## Data and materials availability

The Ku and DNA-PKcs irCLIP data are available via GEO under the accession number <u>GSE109026</u>. PTBP1 and RBFOX2 CLIP data used for controls are available via GEO under accession number <u>GSE78832</u> and <u>ENCODE Data Coordination Center</u> with accession number (ENCSR456FVU). RNA-seq for human primary B cells is available at <u>GSE110669</u>, and for mouse primary B cells are at <u>GSE212194</u>, <u>GSE212195</u>, <u>GSE212196</u>, and <u>GSE214643</u>. The RNA-seq data for *Canis lupus/*dogs and macaque are available at GEO under the accession number <u>GSE125483</u>, for macaque/monkey at <u>GES30352</u>, and for macaque/monkey and Lemuroidea/lemur at <u>Primate Expression Altas</u> and at NHPRTR Tissue Sample Database (SPR021223); for *Equus caballus*/horse are at <u>GSM1139274</u>, <u>GSE43013</u>, and <u>EPR006861</u>. The heatmap and volcano plots were generated using an online plotting tool at https://edb.rdf-lab.org/module/matcalc/.

## Author Information

Correspondence and requests for cellular materials should be addressed to Zha, Shan. Correspondence and requests for irCLIP analyses should be addressed to Zhang, Chaolin

