## Supplementary material for "Ku suppresses RNA-mediated innate immune responses in human cells to accommodate primate-specific Alu expansion": Sup Figure 1-18 and legend

**Extended Data Figure Legends**

**Extended Data Figure 1: Ku, but not LIG4/XRCC4, is required for the viability of human cells and is highly expressed in human cells. a-d,** The Dependency Map analyses of Ku and control genes. *XRCC5/Ku80* (a) and *XRCC6/Ku70* (b) are essential for over 1000 human cancer cell lines tested via CRISPR KO (blue) and RNAi knockdown (purple) screens. Effect score = 0 (neutral, black line) and < -1 (essential, red dashed line) are marked. LIG4 (c) and DNA-PKcs (d) are not commonly essential in human cells tested. **e-g,** The mRNA expression (measured by TPM) for *XRCC5* (e), *XRCC6* (f), and *XRCC4* (g), the obligatory partner of LIG4, in primary human versus mouse B cells. The bars represent means and SEMs and unpaired student's t-test p-value < 0.0001, ****. **h,** The essentiality of Ku in human cells is independent of TERT expression in human cancer cells. Three cancer cell lines (marked by triangles, U2OS, SAOS2, and CAL72) that are known to use ALT to extend their telomere also require Ku for survival. HCT116 is also marked for reference. Data are from the Depmap project.

**Extended Data Figure 2: Generation of Ku80-AID and LIG4-AID tagged cell lines. a,** The strategy for introducing an AID-tag at the N-terminal of endogenous Ku80 or LIG4 with a short linker. The primers used for screens are marked in the diagram and listed in supplementary methods. **b,** The PCR screen of homozygously tagged Ku80 clones 1, 7, and 9 with controls. **c,** The PCR screen of homozygously tagged LIG4 clones 7, 8 and 9 with controls. **d,** Western blotting shows the successful depletion of LIG4 after IAA and Dox treatment. The AID-tagged LIG4 migrated slightly slower than the untagged LIG4. **e,** Western blotting shows that Ku levels remain low for at least 9 days after IAA and Dox treatments. **f,** Representative flow cytometry analyses for the apoptosis marker Annexin V and PI after 6 days of IAA and Dox treatment. **g,** The Annexin V positive cell percentage is plotted at different time points. The bars represent the mean and standard deviation of at least three measurements from two independently derived cell lines. **h**, Representative flow cytometry plot for cell cycle analyses at 48 hr after inducing Ku degradation. S phase percentage was measured by BrdU positivity. The mid-S phase cells were gated, and their BrdU levels were plotted using a histogram. BrdU-negative S phase cells indicate replication stalling. **i**, Quantification of $BrdU^+$/S phase percentage in DMSO versus IAA- and Dox-treated Ku80-AID clones. The bars represent the mean and standard deviation of at least three measurements from two independently derived cell lines. The unpaired student's *t*-test $p < 0.0001$ ****; $p > 0.05$ ns.

**Extended Data Figure 3: Ku degradation induces IFN signaling independent of cycling. a&b,** The gene set enrichment analyses (GSEA) for the interferon (a) and NF-kB (b)-pathways at 4 days after Ku degradation in cycling cells. **c,** The GSEA for the IFN pathway at 24 hr after Ku deletion in cycling cells. **d,** Pathway analyses of significantly downregulated genes upon Ku-degradation (24hr). **e,** Heatmap confirms the G1 arrest after CDK4/6 inhibitor treatment, evidenced by the loss of mitotic and DNA-replication-associated genes (*e.g.*, MCMs and DNA polymerases). The color gradient reflects the Z-scores calculated from normalized TPM across the displayed samples: red represents higher expression levels relative to the mean, while blue indicates lower expression levels. **f &g,** The GSEA for the interferon-pathway at 48hr (2 days, f) and 96 hr (4 days, g) after Ku deletion in G1 arrested cells. **h,** The GSEA for NF-kB pathway at 4 days after Ku-degradation in G1 arrested cells. **i, j and k,** Quantitative RT-PCR analyses of NF-kB target genes (GADD45B(i); Jun(j); and PPP1R15A(k)). Each dot represents a biological replication; three Ku80-AID clones (#1, 7, 9) and three HEK293 clones with inducible Ku (Sal, SB, and T1) were analyzed individually and plotted together. The mRNA levels were normalized against β-actin. **l**, Quantification of NF-kb p50/p105 ratio from multiple independent western blots on independent clones. The unpaired student's *t*-test $p<0.0001$ ****.

**Extended Data Figure 4: Ku deletion in HCT116 also induces IFN signaling. a** PCR analyses show the specific and efficient deletion of the Ku86 flox allele upon 4OHT treatment. **b**, Progressive loss of viability in 4OHT treated *Ku86$^{Flox/-}$:Cre-ERT2*, but not control *Ku86$^{Flox/-}$* cells. **c,** Pathway analyses of significantly up-regulated genes upon Ku-deletion (96 hr). **d &e,** The GSEA for the interferon-pathway at 96 hr (4 days) after 4OHT treatment in control *Ku86$^{flox/-}$* cells (d) or *Ku86$^{flox/-}$:Cre-ERT2* (e). **f,** Heatmap of selective NF-kB targets in Ku-AID and Ku-flox cells. The color gradient reflects the Z-scores calculated from normalized TPM across the displayed samples: red represents higher expression levels relative to the mean, while blue indicates lower expression levels.

**Extended Data Figure 5: MAVS but not c-GAS-STING mediates IFN activation upon Ku degradation a,** Western blotting shows that cGAS protein is not expressed in HCT116 cells with Hela cells as the positive control. **b,** Ku is required in human cells regardless of cGAS expression levels**.** The mRNA levels of cGAS, a dsDNA sensor provided by the Dependency Map database, were plotted against the *XRCC5/Ku80* Gene deletion effect. Effect score =0 (neutral) and <-1 (essential). **c-g,** Quantitative RT-PCR analyses of IFN targets, ISG15 (**c**), IFIT1(**d**), IFNb(**e**), STAT1 (**f**), and NF-kB target GADD45B (**g**) validated the critical role of MAVS in mediating IFN activation following Ku-degradation. Three independently derived Ku80-AID clones and three independent MAVS KO clones were analyzed individually and plotted together. PolyI:C transfection (100 ng/ml, 24 hr) was used as a positive control. **h,** Representative western blotting analyses of IFN pathway markers in multiple independently derived MAVS, RIG-I, and MDA5 KO clones. Additional clones can be found in **Fig. 3a**. **i,** Western blotting analyses of dsRNA sensors RIG-I, MDA5, MAVS, and IRF7 in HCT116, HEK293 and control Hela cells.

**Extended Data Figure 6: Ku depletion induces PKR and OAS activation beyond IFN. a,** Purified IFNβ (up to 40 ng/ml) causes at most 10% decrease of total cellularity in three days. **b,** Transfected poly IC of 10 ng/ml and 100 ng/ml causes 40% and 99% lost of viability in 3 days regardless of Ku-AID tag (no IAA or Dox). Multiple repeats with consistent results were analyzed and one set of representative experiments was plotted. The bars represent the mean and SEM from multiple biological repeats. **c**, Relative increase of OP-puro low population in Ku80-AID cells after IAA and Dox treatment. We note that Dox also causes a mild decrease of OP-puro in a small subset of parental AID cells. Dox, a tetracycline antibiotic, is known to inhibit bacterial protein synthesis by binding to the 30S ribosomal subunit. While it is highly selective for prokaryotes, Dox can mildly suppress protein translation in human cells by impairing mitochondrial translation[1]. To ensure the translation effect was caused by Ku depletion, not Dox or IAA, we also measured the OP-puro levels in 4OHT treated *Ku86$^{flox/-}$:CreERT2* and control *Ku86$^{flox/-}$*cells with minimal background translation changes **(Fig. 3c)**. **d**, Multiple members of the OAS family are transcriptionally induced (up to 128-fold for OAS1) upon Ku degradation (day 4 after Ku degradation). The error bars represent the standard errors from three independently derived cell lines. **e,** The 28S versus 5S and 5.8S rRNA ratios in control DMSO-treated cells versus IAA- and Dox-treated Ku80-AID cells, and poly I:C (100 ng/ml) transfected cells. **f,** The representative histogram shows the relative reduction of 28S size and the relative decrease of 28S rRNA abundance after Ku80 degradation. **g,** The 28S versus 18S rRNA ratio in 4OHT treated control *Ku86*$^{flox/-}$ cells and *Ku86*$^{flox/-}$*:CreERT2* cells (Day7). For panel c, d, e, and g, at least three independent samples were measured at each time point from multiple clones. The means and SEMs were plotted. The *p*-values were calculated via an unpaired student *t*-test, p<0.05,*;<0.01,**; <0.001, ***; and <0.0001, ****.

**Extended Data Figure 7: Ku deletion in HEK293 cells induces NF-kB signaling and PKR and RNaseL activation. a,** Dox (1 µg/ml) induces ~10 fold Ku70 mRNA overexpression the engineered HEK293 cells. CRISPR KO only removed a small exon fragment from endogenous Ku70, therefore, even after Dox is withdrawn, RNA-seq can still detect endogenous Ku70 mRNA (although it is not able to encode Ku70 protein). **b,** Western blot shows the slow loss of Ku, the robust NF-kB and PKR activation, in HEK293 cells after Dox withdrawal. **c**, The moderate and significant loss of viability upon Dox withdrawal in HEK293 cells. **d** Pathway analyses of significantly upregulated genes upon Ku-depletion in HEK293 cells (8 days). **e,** The GSEA for the NF-kB-pathway at 8 days after Dox was withdrawn. Three independently derived clones were analyzed and plotted together. **f,** The HEK293 cells do not express IRF7 mRNA. See **Extended Data Fig. 5i** for data support that IRF7 protein also can not be induced in the HEK293 after IFN treatment. **g**, Gating strategy and quantification of protein translation by OP-Puro in HEK293 cells after Dox withdrawal at 6 days, n=3 clones with 3 experimental repeats. **h,** The 28S versus 5S and 5.8S rRNA ratio in Dox-treated and Dox withdrawal (8 days) HEK293 cells with inducible Ku70 expression.

**Extended Data Figure 8: CRISPR KO of MAVS, RIG-I, and PKR and higher primates have increased Ku expression.**
**a,** The validation of PKR CRISPR KO. PKR is an IFN inducible gene. The poly I:C treatment induced maximal PKR expression and activation only in parental cells and ensured a clean CRISPR KO**. b,** Western blotting analyses of Ku in human, non-human primates, and non-primate cell lines. **c-e,** The mRNA expression levels of XRCC5(c), XRCC6(d), and ADAR1(e) were analyzed across species. Human and mouse data came from our previous studies, while other species' data were from public datasets. **f-h,** AlphaFold3 modeling of core human Ku (XRCC5 aa 1–536, pink/purple; XRCC6 aa 36–521, yellow/orange) bound to asSL1 RNA of Alu (70nt, silver) or perfectly paired dsDNA (58nt, gold). In each model, light-shaded proteins (pink and yellow) represent Ku bound to asSL1 RNA, while darker-shaded proteins (purple and orange) represent Ku bound to dsDNA. The top view **(f)** and front-side view **(g)** are presented in overlay format. The inset shows that base U19 and U21 of asSL1 flip out in the Ku-bound model but not in the absence of Ku. **Panel (h)** compares the structures of Ku-bound dsDNA (top, Ku hidden), Ku-bound asSL1 RNA (middle, Ku hidden), and free asSL1 RNA. The hairpin diagram was generated using the IDT oligo analysis tool. The U19 and U21 bases of asSL1 flip out to accommodate Ku binding. See Methods for further details.

**Extended Data Figure 9: Ku binds to SINEs without selective preference for a specific Alu subfamily. a-b,** Two other nuclear RBPs are shown to compare to Ku and DNA-PKcs, highlighting the strong preference for introns and repetitive elements for Ku and DNA-PKcs. **c-d,** The genomic distribution of DNA-PKcs irCLIP tags (c) and overlap with repetitive elements (d)**. e-f, as**Alu elements of specific subfamilies with the numbers overlapping with Ku (e) and DNA-PKcs (f) irCLIP peaks shown in the y-axis and the total numbers in the human genome shown in the x-axis. The red arrows indicate Alu subfamilies with lower-than-expected irCLIP signal.

**Extended Data Figure 10: Ku binds to monomeric Alu at the stem loops. a-b,** The relative fraction of irAlu and non-irAlu defined by Wu et al. *Cell 2018* [2] bound by Ku (a) or DNA-PKcs (b). **c,** The distribution of LINE element orientation among the Ku and DNA-PKcs irCLIP sites. **d,** The irCLIP sites preference of Ku in asAlu elements. The red is a higher frequency. **e,** The irCLIP sites preference of DNA-PKcs in asAlu elements. The red is a higher frequency **f,** The impact of irCLIP-Ku on mRNA expression. The Log2 (mean TPM+1) in DMSO versus IAA- and Dox- treated (day 4) Ku80-AID samples were plotted. The red and blue dots are genes with irCLIP-Ku in 3'-UTR (with or without intron, red), or in intron only (blue) **g,** The impact of Alu on mRNA expression after Ku degradation. The Log2 (mean TPM+1) in DMSO versus IAA- and Dox-treated (day 4) Ku80-AID samples were plotted. The red and blue dots are genes with annotated Alus in 3'-UTR (with or without Intron, red, n>2), or in Intron only (blue, n>9). The data associated with panels f and g can be found in **Extended Data Tables 12 and 13**.

**Extended Data Figure 11: The irCLIP sites for Ku and DNA-PKcs are enriched at the stem-loop regions of asAlu. (a-c),** DNA-PKcs**. (d-f),** Ku. Ku and DNA-PKcs binding sites are represented by irCLIP peaks (a, d), or crosslink sites inferred by CIMS (b, e) or CITS (c, f) analysis. Each panel shows the fraction of Ku or DNA-Pkcs binding sites mapped to each position of asAlu normalized by the total number of binding sites.

**Extended Data Figure 12: Representative Ku and DNA-PKcs irCLIP binding sites in *MDM4* gene.** *MDM4* has several irCLIP peaks and crosslink sites (CIMS and CITS) for Ku and DNA-PKcs in the 3'-UTR, including several sites overlapping with Alu elements. One site in the 3'-UTR overlapping with antisense Alu is highlighted with a zoomed-in view.

**Extended Data Figure 13: Representative Ku and DNA-PKcs irCLIP binding sites in *LOC284454* and *Mir23*.** Ku and DNA-PKcs peaks and crosslink sites (CIMS and CITS) overlapping with *Mir23* are highlighted in the zoom-in view at the top. Ku and DNA-PKcs binding sites overlapping with an asAlu in *LOC284454* are highlighted in the zoomed-in view at the bottom.

**Extended Data Figure 14: Ku binds individual Alu SL with high affinity. a,** Schematic illustration of the Ku Fluorescence Polarization (FP) assay. FAM-labeled ligands are incubated with increasing concentrations of Ku protein. The FAM-fluorophore is excited with linearly polarized light. Rapidly tumbling free FAM-ligand emits depolarized light, while the Ku/FAM-ligand complex tumbles slower, leading to the emission of relatively more polarized light. **b,** SDS-gel of purified Ku heterodimer used in the FP assay. Protein standard: BioRad Precision Plus Protein™ Kaleidoscope™. **c,** Schematic representation of the Alu consensus secondary structure model with SL1 and SL2. Ligands used in this study are a dsDNA stem-loop, the Alu SL1 and the antisense SL1 (asSL1) with Vienna RNAfold predicted secondary structures[3]. **d,** FP binding curves of dsDNA, SL1 and asSL1 dsRNA with increasing concentrations of Ku. Errors are given as standard deviation from replicates. $K_D$ values are given in the inset.

**Extended Data Figure 15: Correlation between Ku-irCLIP peaks versus DNA-PKcs irCLIP peaks on Alu.** The number of irCLIP tags by Ku was plotted against the number irCLIP tags by DNA-PKcs. The sense (blue) and antisense (grey) Alus were plotted separately. The data were derived from numbers in **Extended Data Table 9**.

**Extended Data Figure 16: Ku degradation does not significantly increase dsRNA detected by the J2 antibody. a-b,** Representative J2 antibody staining (a) and quantification (b) before and after IAA- and Dox-induced Ku80 degradation. **c,** Representative J2 staining from human HCT116 cells versus immortalized murine embryonic fibroblasts (iMEF). Two independent fields were shown for both cell lines, labeled as view 1 and view 2. **d,** Quantification of J2 antibody staining in HCT116 cells and iMEFs. **e,** Representative J2 staining in nuclease treated cells. DNaseQ1 cleaves DNA, RNAase T1 cleaves ssRNA, and RNase III cleaves dsRNA.

**Extended Data Figure 17: Representative Ku and DNA-PKcs irCLIP binding sites in *tRNA-Lys-AAA and tRNA-Cys-TGY loci.*** Ku and DNA-PKcs peaks and crosslink sites (CIMS and CITS) overlapping with *tRNA loci* are highlighted in the zoom-in view at the top. Ku and DNA-PKcs binding sites overlapping with tRNA independent of Alu.

**Extended Data Figure 18: The irCLIP sites for Ku and DNA-PKcs are enriched at the anti-coden loop and the TΨC loop of t-RNA. (a-b),** DNA-PKcs**. (c-d),** Ku**.** Ku and DNA-PKcs crosslink sites inferred by CIMS (**a, c**) or CITS (**b, d**) analysis. Each panel shows the fraction of Ku or DNA-Pkcs binding sites mapped to each position of tRNA normalized by the total number of binding sites. **e,** A diagram shows the representive tRNA folding. The three major loop regions are also highlighted in (b, d). The sequence is derived from a yeast Phe tRNA (adapted from https://en.wikipedia.org/wiki/Transfer_RNA).

### Extended Data Figure 1

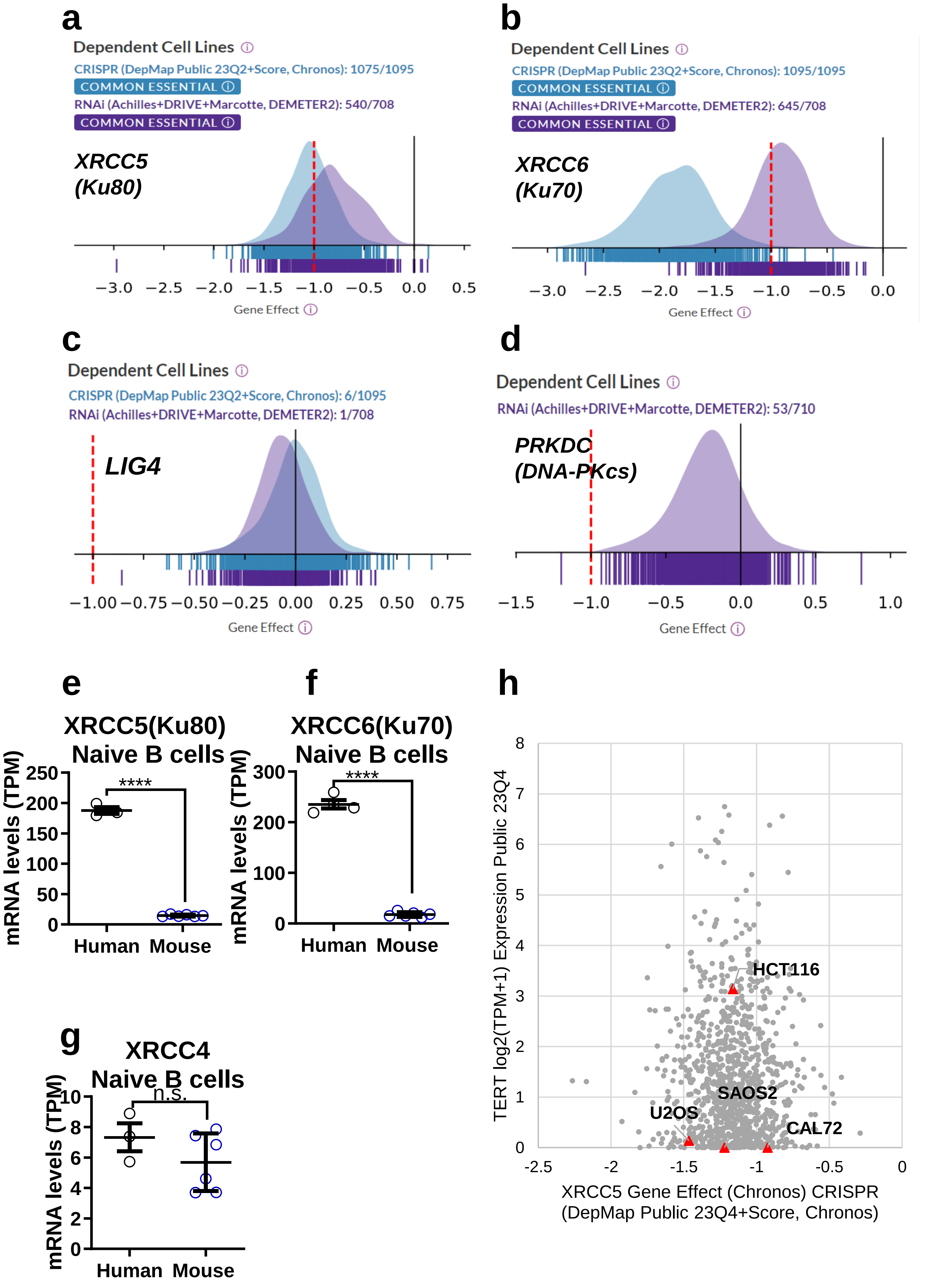

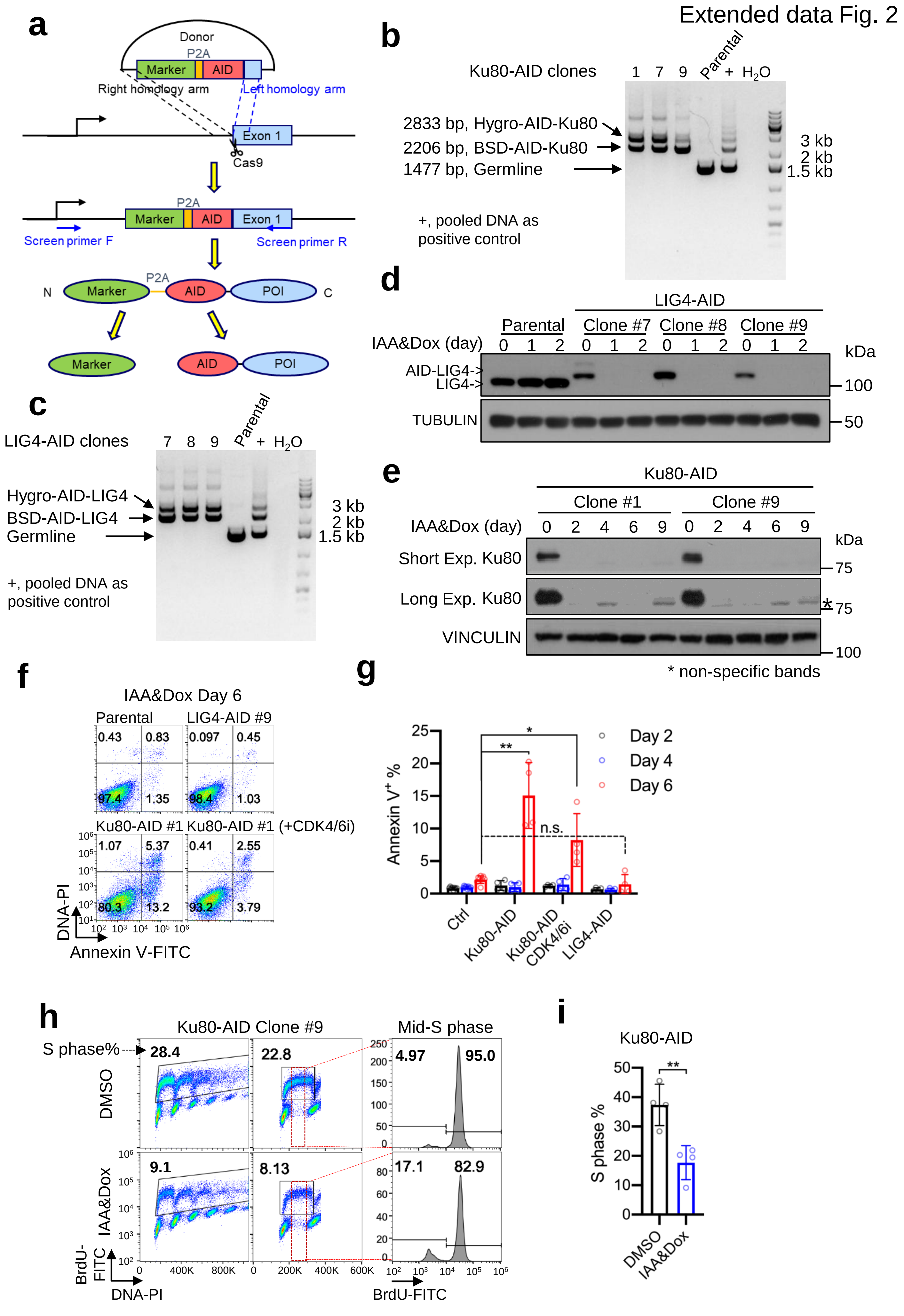

Extended data Fig. 2
a
Donor
P2A
Marker
AID
Right homology arm
Left homology arm
Exon 1
Cas9
Screen primer F
Screen primer R
N
POI
C
b
Ku80-AID clones
1 7 9 Parental + $H_2O$
2833 bp, Hygro-AID-Ku80
2206 bp, BSD-AID-Ku80
1477 bp, Germline
+, pooled DNA as positive control
3 kb
2 kb
1.5 kb
c
LIG4-AID clones
7 8 9 Parental + $H_2O$
Hygro-AID-LIG4
BSD-AID-LIG4
Germline
+, pooled DNA as positive control
d
LIG4-AID
Parental
Clone #7
Clone #8
Clone #9
IAA&Dox (day)
0 1 2 0 1 2 0 1 2 0 1 2
AID-LIG4->
LIG4->
TUBULIN
kDa
100
50
e
Ku80-AID
Clone #1
Clone #9
IAA&Dox (day)
0 2 4 6 9 0 2 4 6 9
Short Exp. Ku80
Long Exp. Ku80
VINCULIN
kDa
75
75
100
* non-specific bands
f
IAA&Dox Day 6
Parental
LIG4-AID #9
Ku80-AID #1
Ku80-AID #1 (+CDK4/6i)
0.43 0.83 97.4 1.35
0.097 0.45 98.4 1.03
1.07 5.37 80.3 13.2
0.41 2.55 93.2 3.79
DNA-PI
Annexin V-FITC
g
Annexin $V^+$ %
Day 2
Day 4
Day 6
**
*
n.s.
Ctrl
Ku80-AID
Ku80-AID CDK4/6i
LIG4-AID
h
Ku80-AID Clone #9
Mid-S phase
S phase%
DMSO
IAA&Dox
28.4
22.8
4.97 95.0
9.1
8.13
17.1 82.9
BrdU-FITC
DNA-PI
i
Ku80-AID
S phase %
**
DMSO
IAA&Dox

Extended data Fig. 3

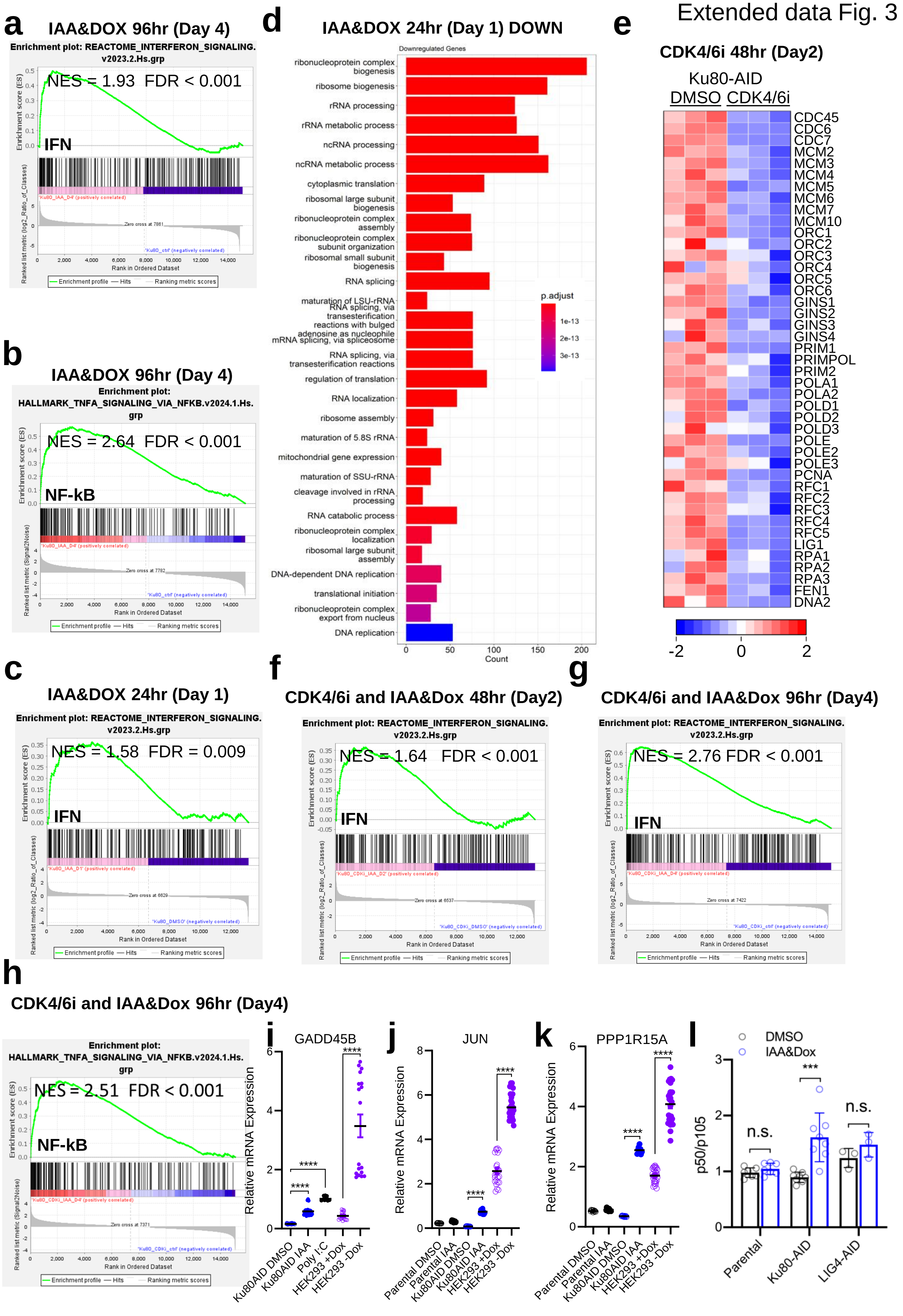

**a**

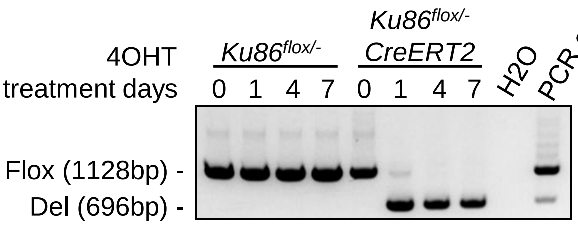


**b**

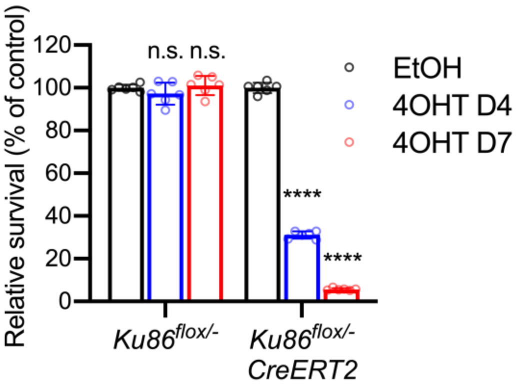


**c** ***Ku86$^{flox/-}$:CreERT2*** **+/- 4OHT 96hr Deseq2 FDR<0.01 and FC>2**

| Group | Pathway | # Genes in overlap (k) | p | %hits |
|---|---|---|---|---|
| UP | **INTERFERON_ALPHA_RESPONSE** | 26 | 0.00E+00 | 5.71% |
| UP | **INTERFERON_GAMMA_RESPONSE** | 32 | 1.04E-10 | 7.03% |
| UP | **TNFA_SIGNALING_VIA_NFKB** | 20 | 1.04E-10 | 4.40% |
| UP | **P53_PATHWAY** | 32 | 1.04E-10 | 7.03% |
| UP | INFLAMMATORY_RESPONSE | 17 | 1.74E-10 | 3.74% |
| UP | APOPTOSIS | 15 | 3.15E-10 | 3.30% |
| UP | HYPOXIA | 12 | 1.92E-06 | 2.64% |
| UP | DNA_REPAIR | 6 | 5.67E-03 | 1.32% |
| UP | IL2_STAT5_SIGNALING | 7 | 5.81E-03 | 1.54% |
| UP | IL6_JAK_STAT3_SIGNALING | 4 | 1.43E-02 | 0.88% |

**d** ***Ku86$^{flox/-}$*** **+/- 4OHT 96hr**

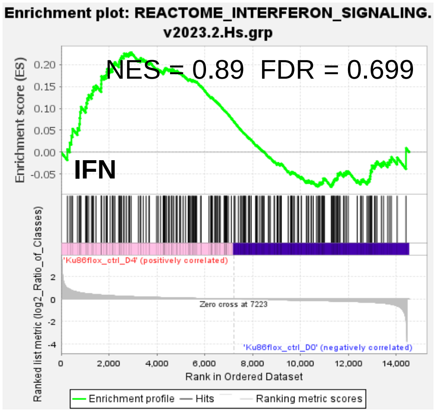


**e** ***Ku86$^{flox/-}$:CreERT2*** **+/- 4OHT 96hr**

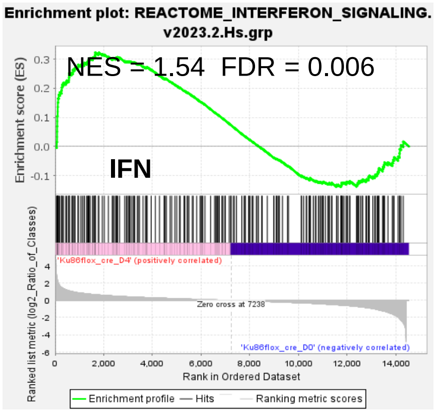


**f**

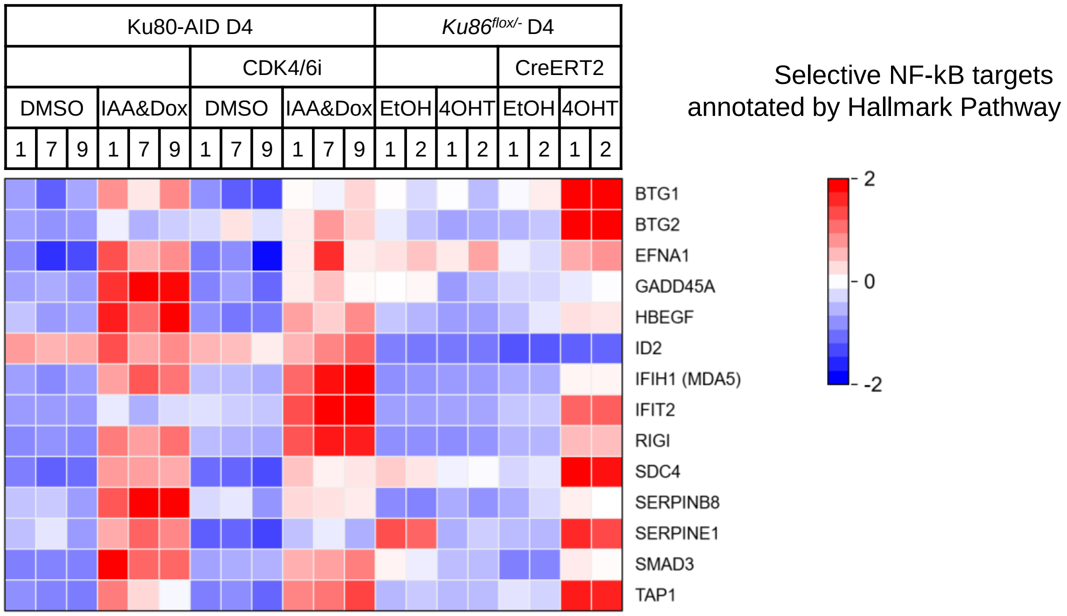

Extended data Fig. 5

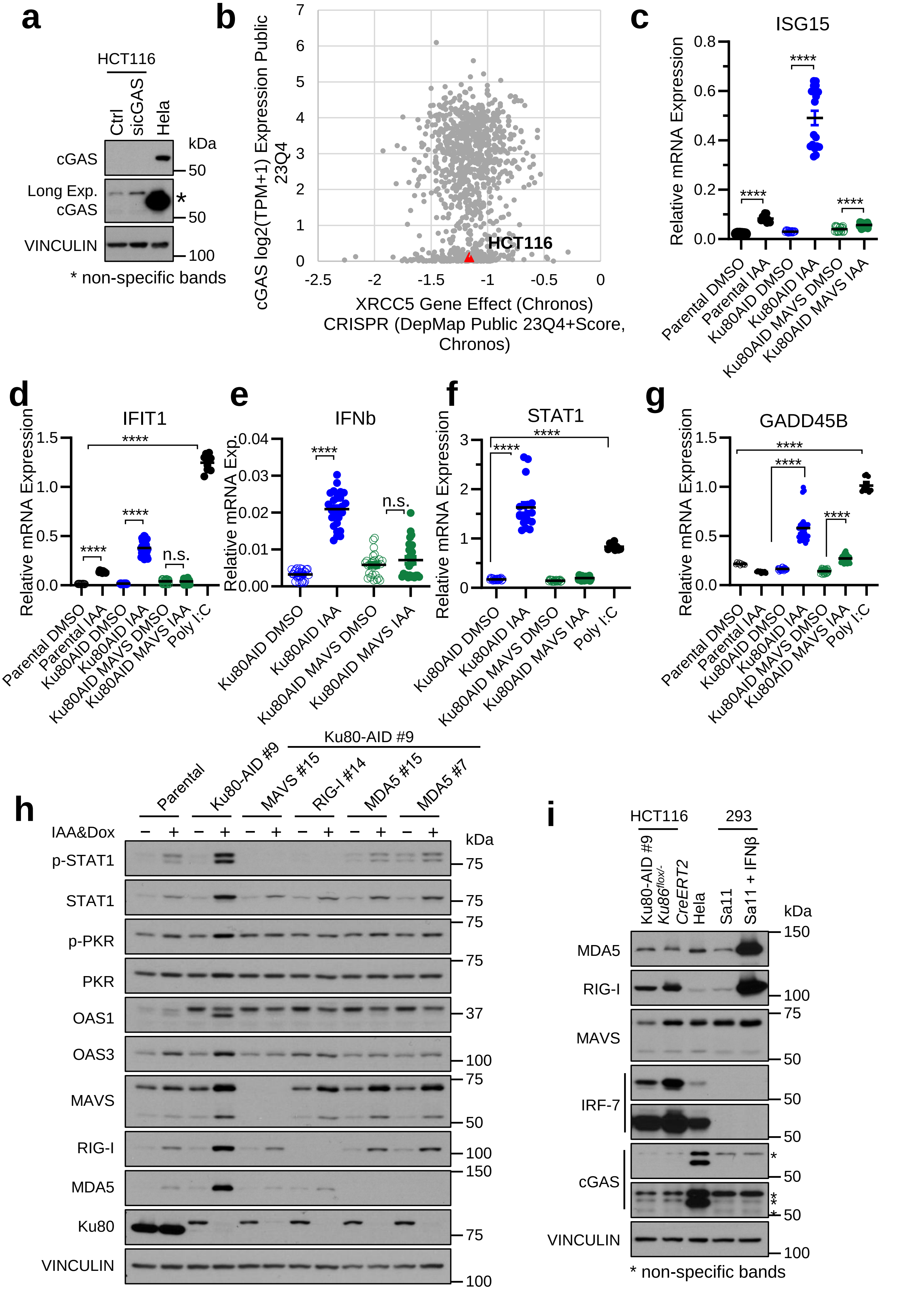

Extended data Fig. 6

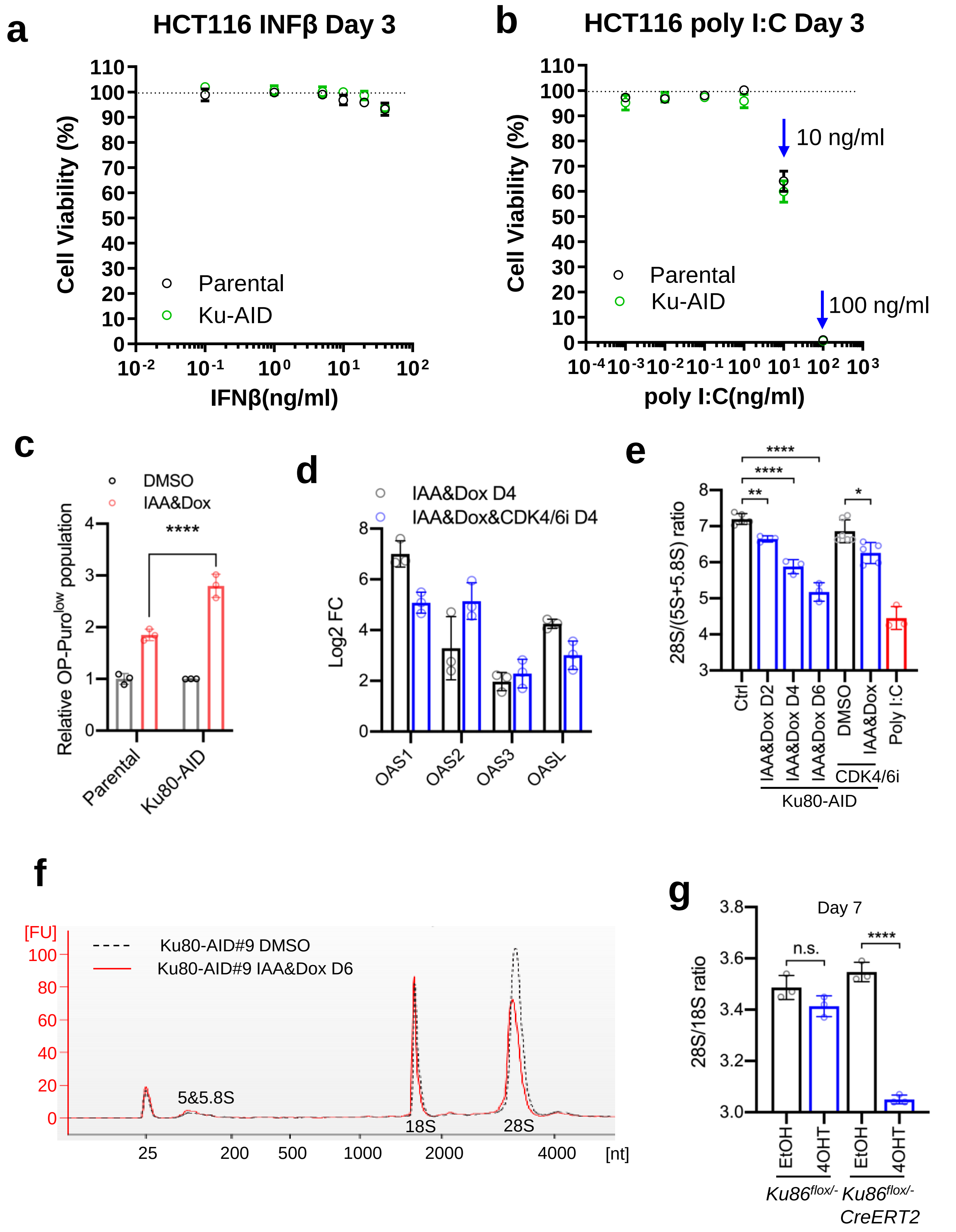

Extended data Fig.7

a

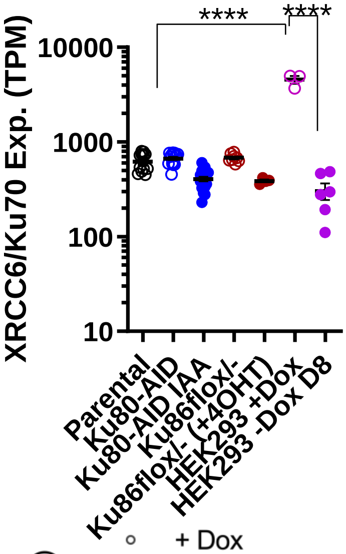


b

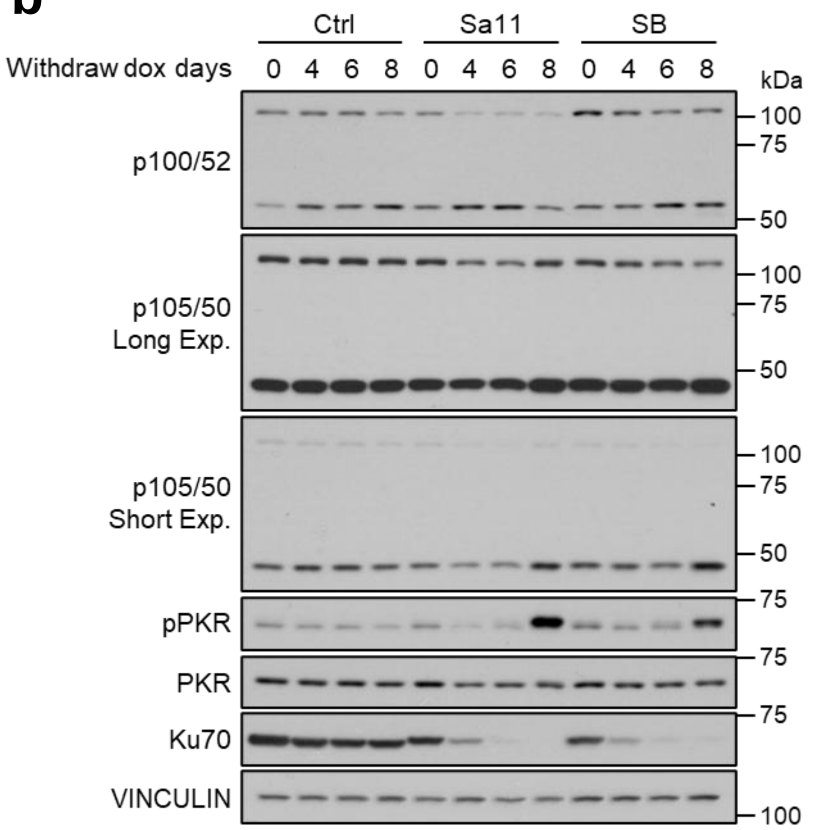


c

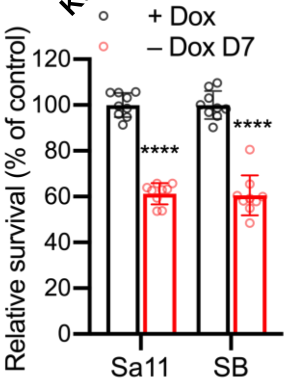


d

| | Pathway | # Genes in overlap (k) | p | %hits |
|---|---|---|---|---|
| UP | **TNFA_SIGNALING_VIA_NFKB** | 48 | 0.00E+00 | 3.96% |
| UP | UNFOLDED_PROTEIN_RESPONSE | 23 | 0.00E+00 | 1.90% |
| UP | P53_PATHWAY | 33 | 0.00E+00 | 2.72% |
| UP | UV_RESPONSE_UP | 21 | 5.02E-09 | 1.73% |
| UP | DNA_REPAIR | 19 | 6.09E-08 | 1.57% |
| UP | APOPTOSIS | 18 | 8.85E-07 | 1.49% |
| UP | **INTERFERON_ALPHA_RESPONSE** | 7 | 2.09E-02 | 0.58% |
| UP | IL2_STAT5_SIGNALING | 11 | 2.80E-02 | 0.91% |
| UP | **INTERFERON_GAMMA_RESPONSE** | 11 | 2.88E-02 | 0.91% |
| UP | IL6_JAK_STAT3_SIGNALING | 6 | 3.79E-02 | 0.50% |
| UP | UV_RESPONSE_DN | 8 | 5.43E-02 | 0.66% |
| UP | INFLAMMATORY_RESPONSE | 10 | 6.12E-02 | 0.83% |

e

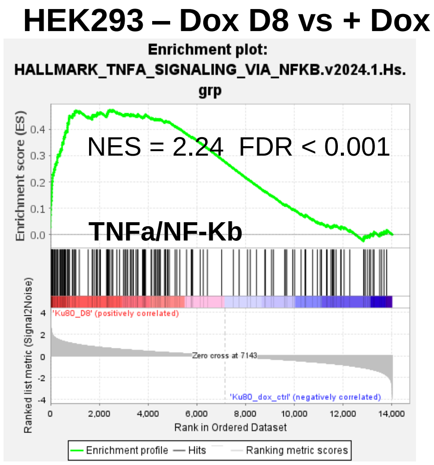


f

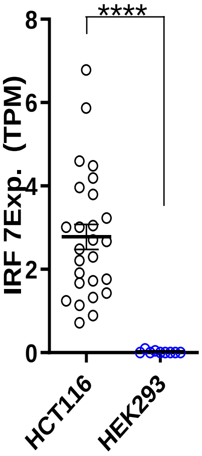


g

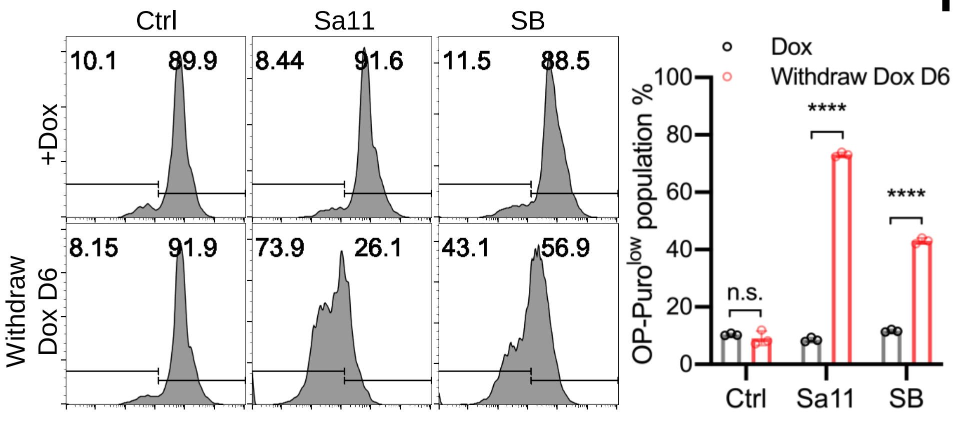


h

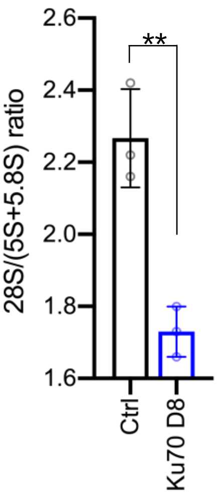

Extended data Fig. 8

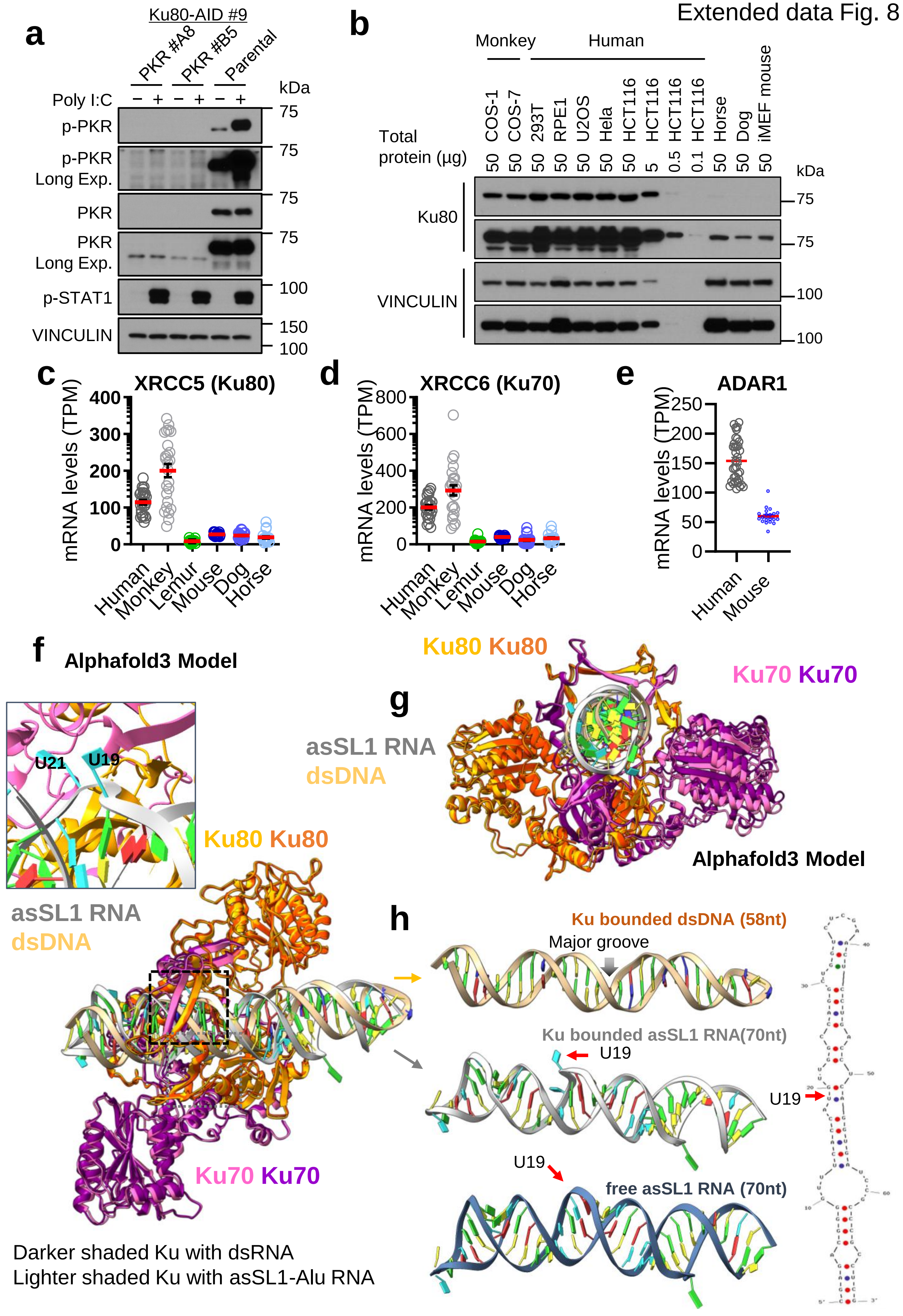

Extended data Fig. 9

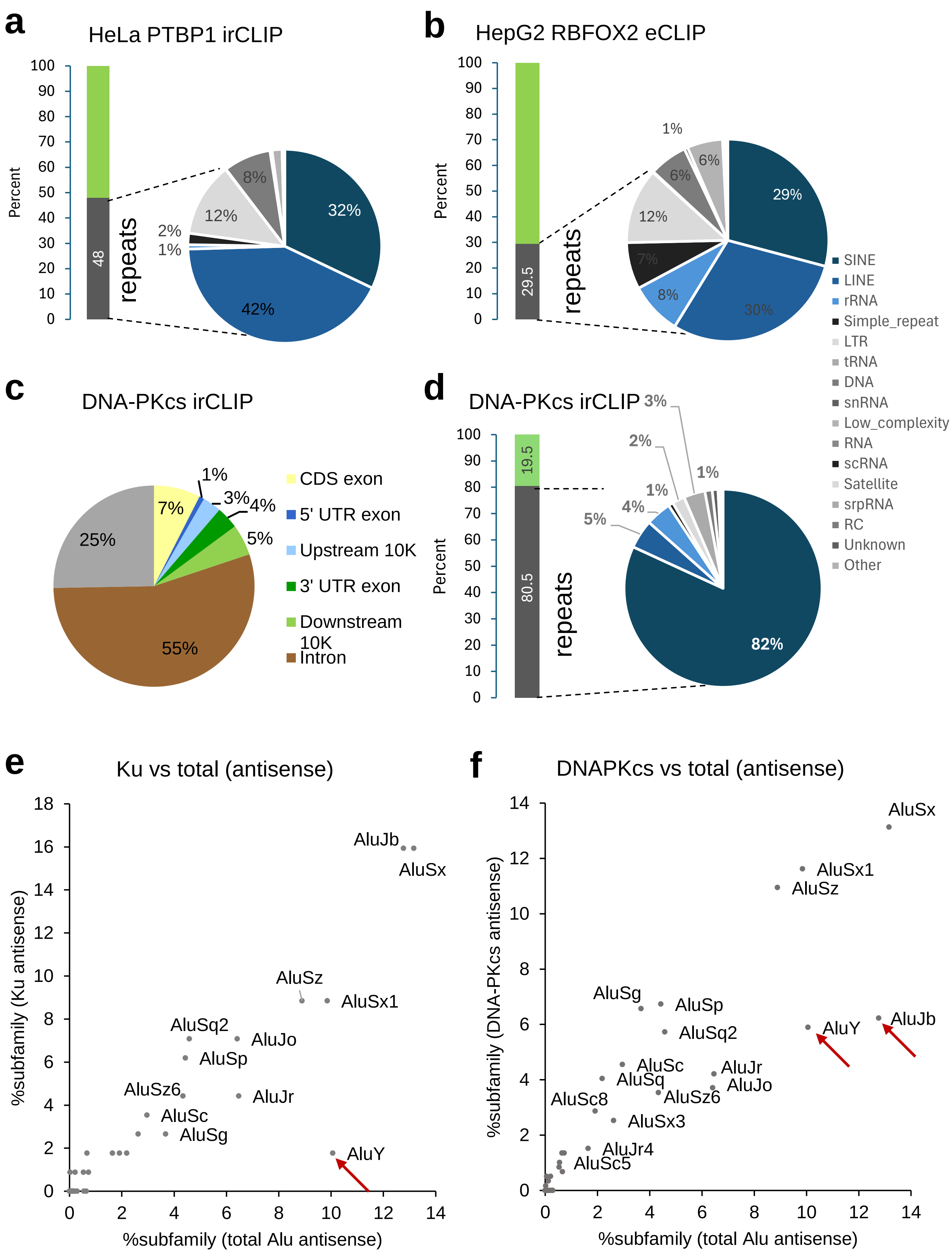

Extended data Fig. 10

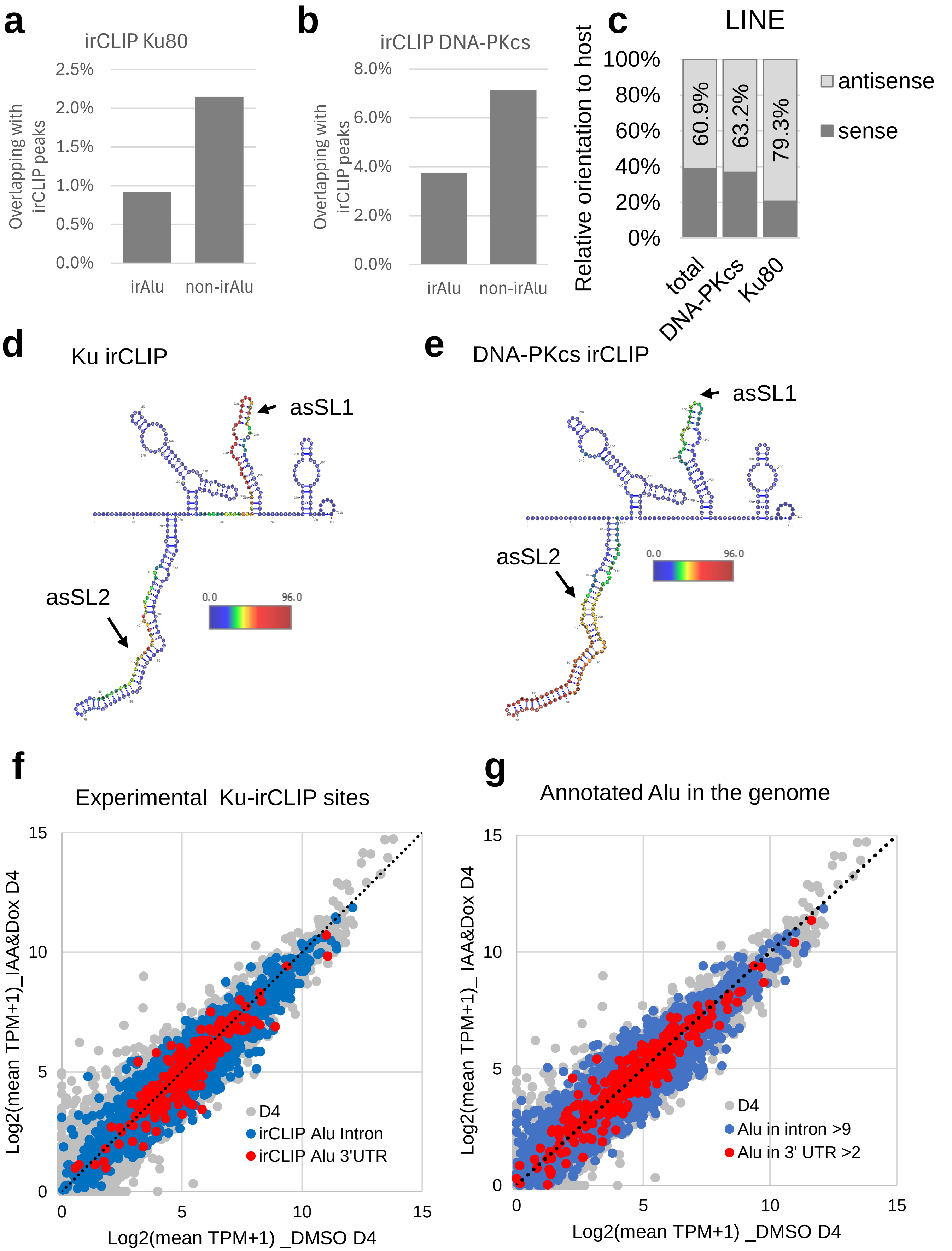

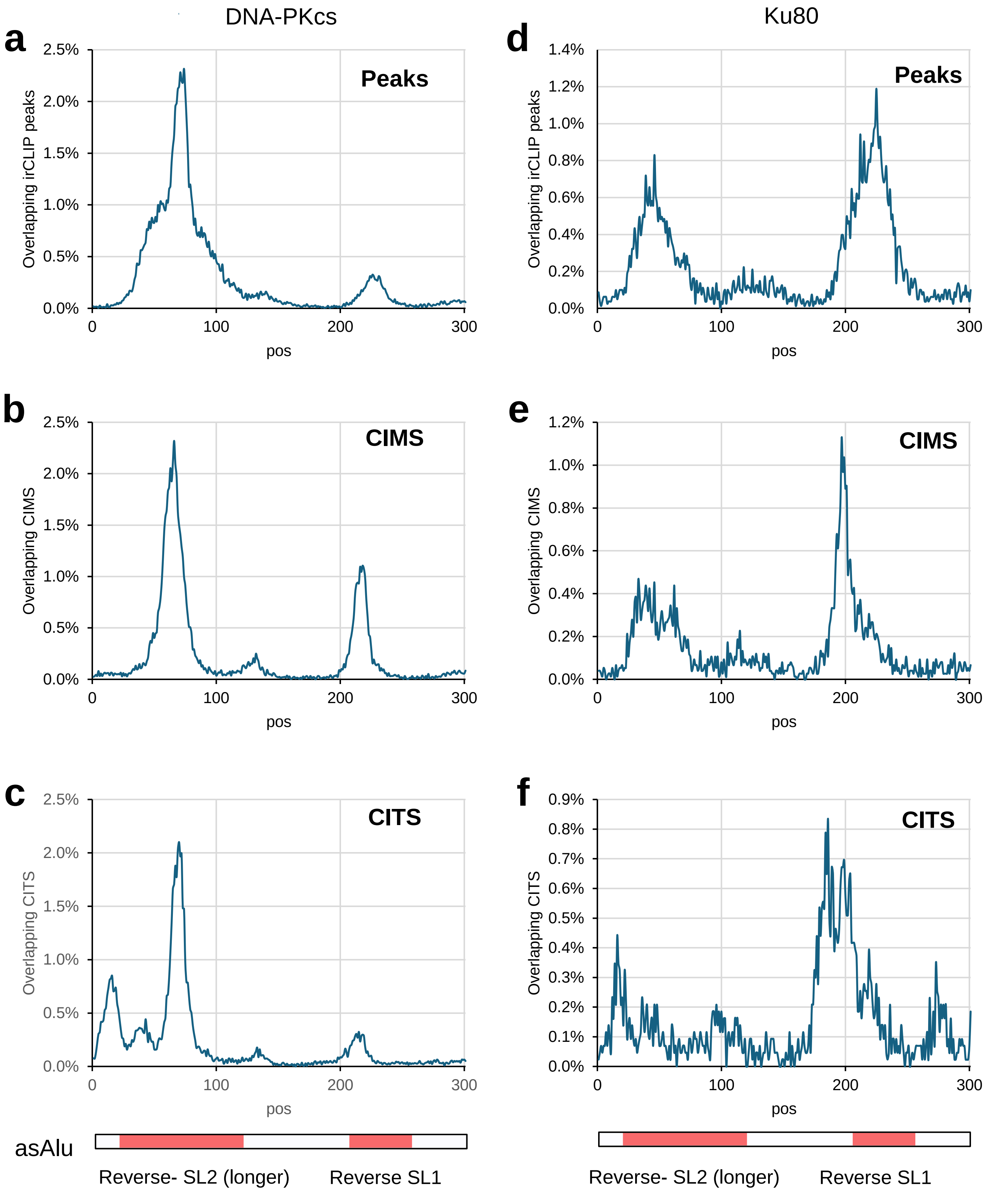
DNA-PKcs
Ku80
a
Peaks
Overlapping irCLIP peaks
2.5%
2.0%
1.5%
1.0%
0.5%
0.0%
0
100
200
300
pos
d
1.4%
1.2%
1.0%
0.8%
0.6%
0.4%
0.2%
0.0%
b
CIMS
Overlapping CIMS
e
c
CITS
Overlapping CITS
f
0.9%
0.7%
0.5%
0.3%
0.1%
asAlu
Reverse- SL2 (longer)
Reverse SL1

Extended data Fig. 12

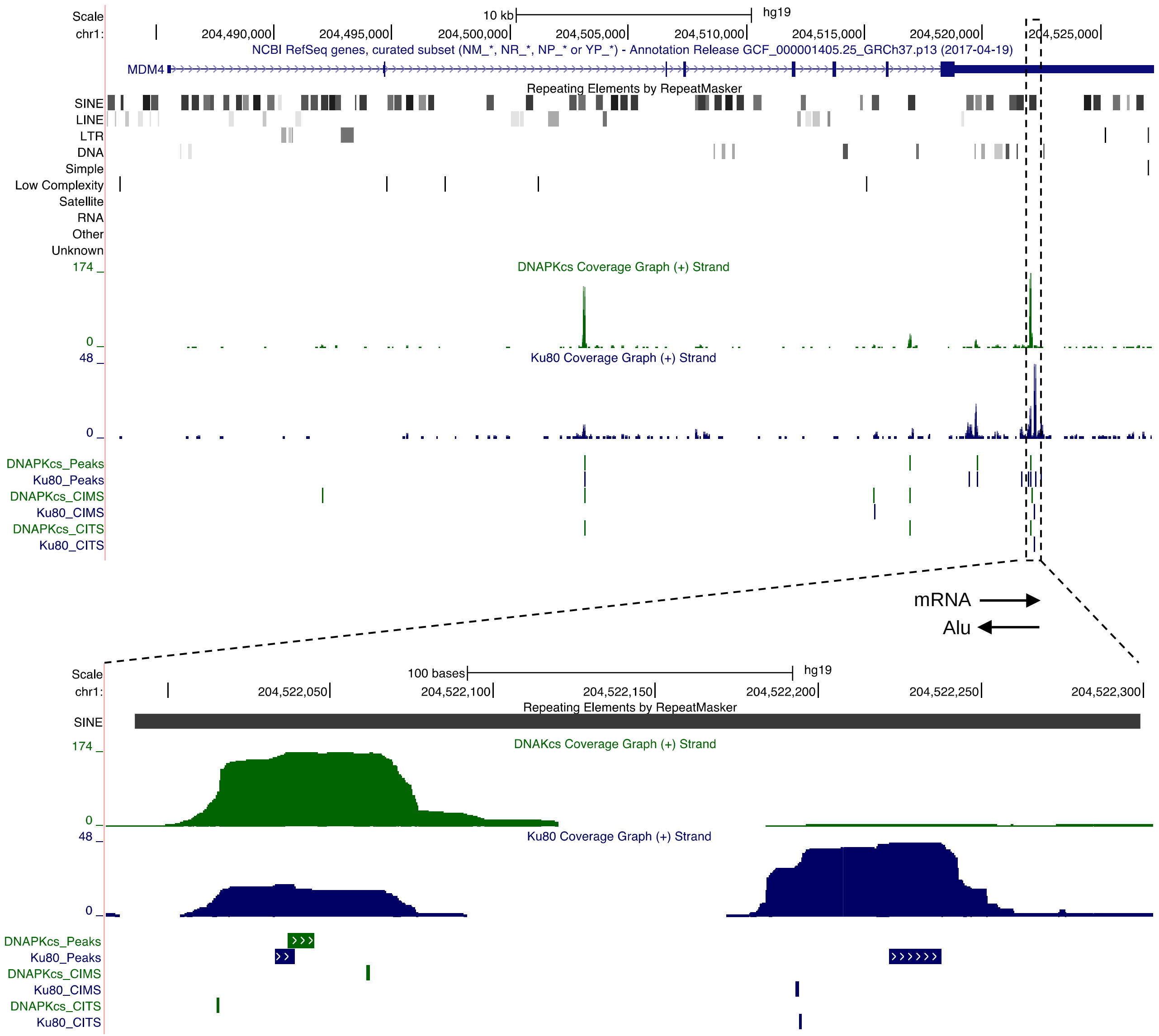

### Extended data Fig. 13

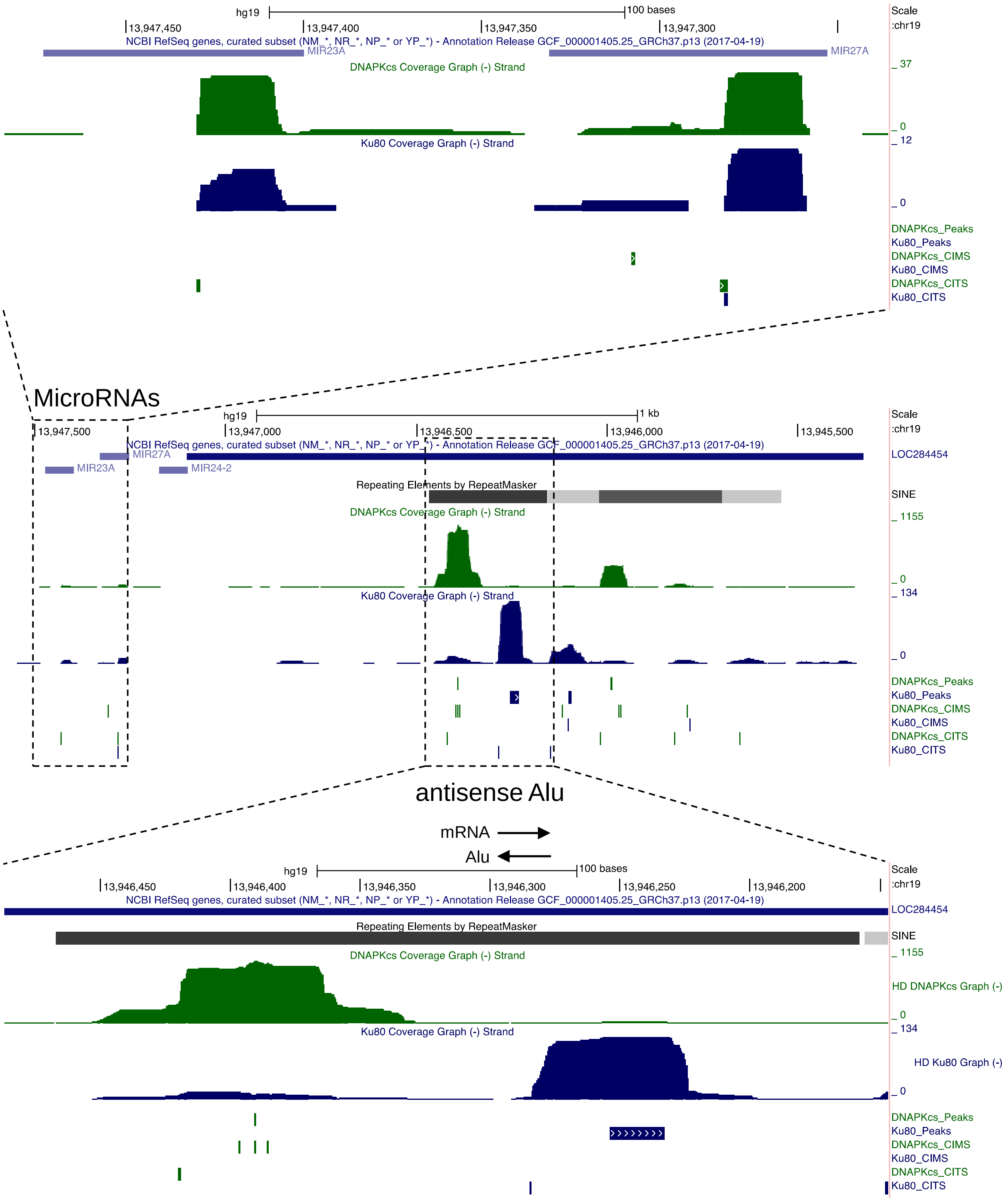

Extended data Fig. 14

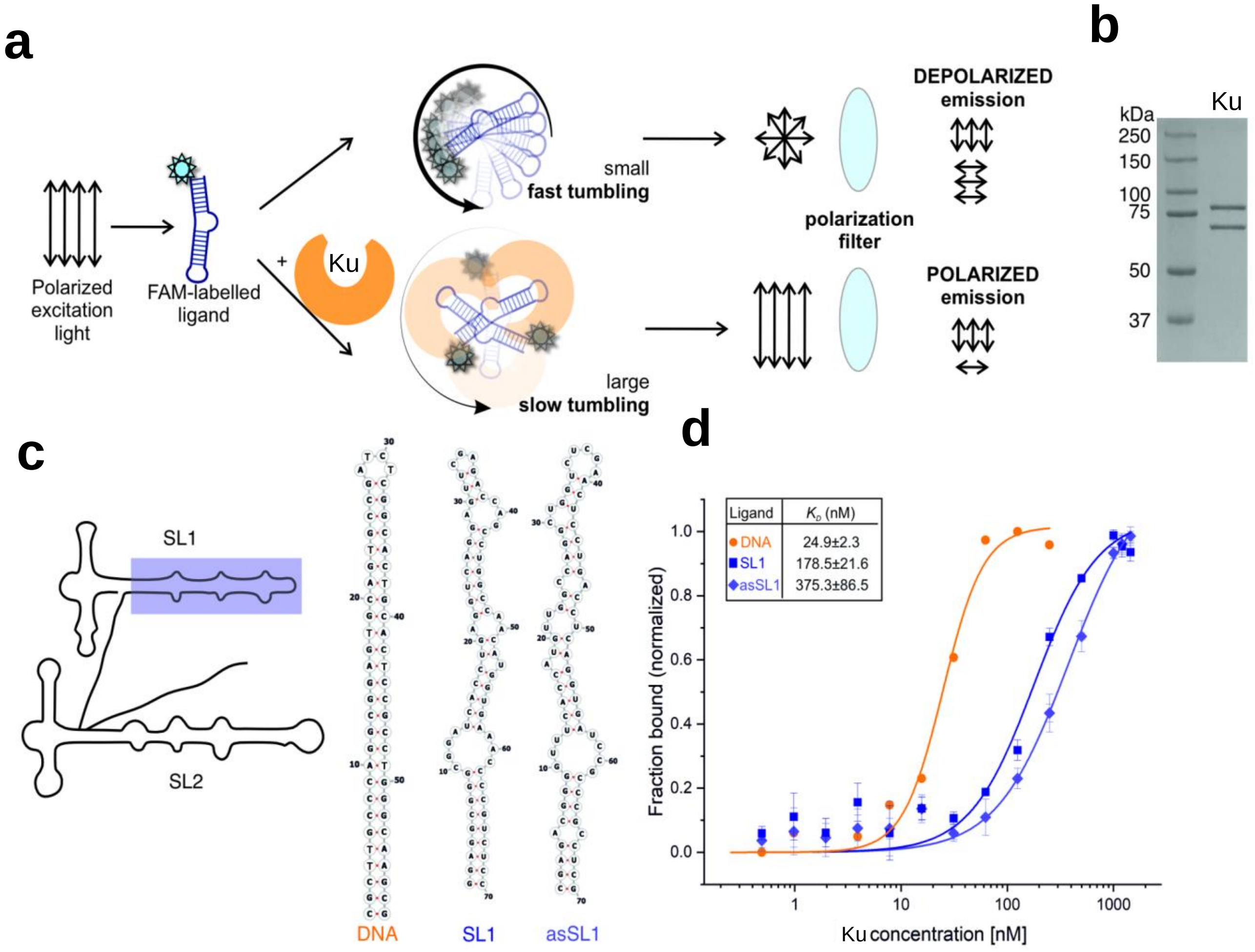

Extended data Fig. 15

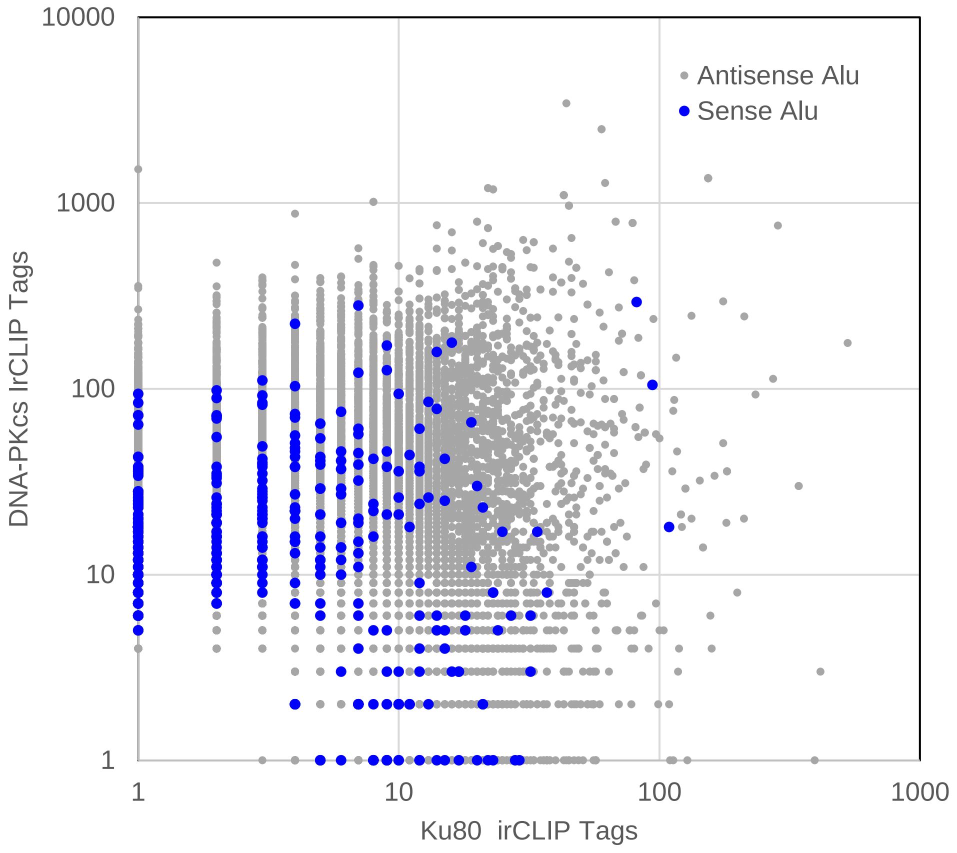

Extended data Fig. 16

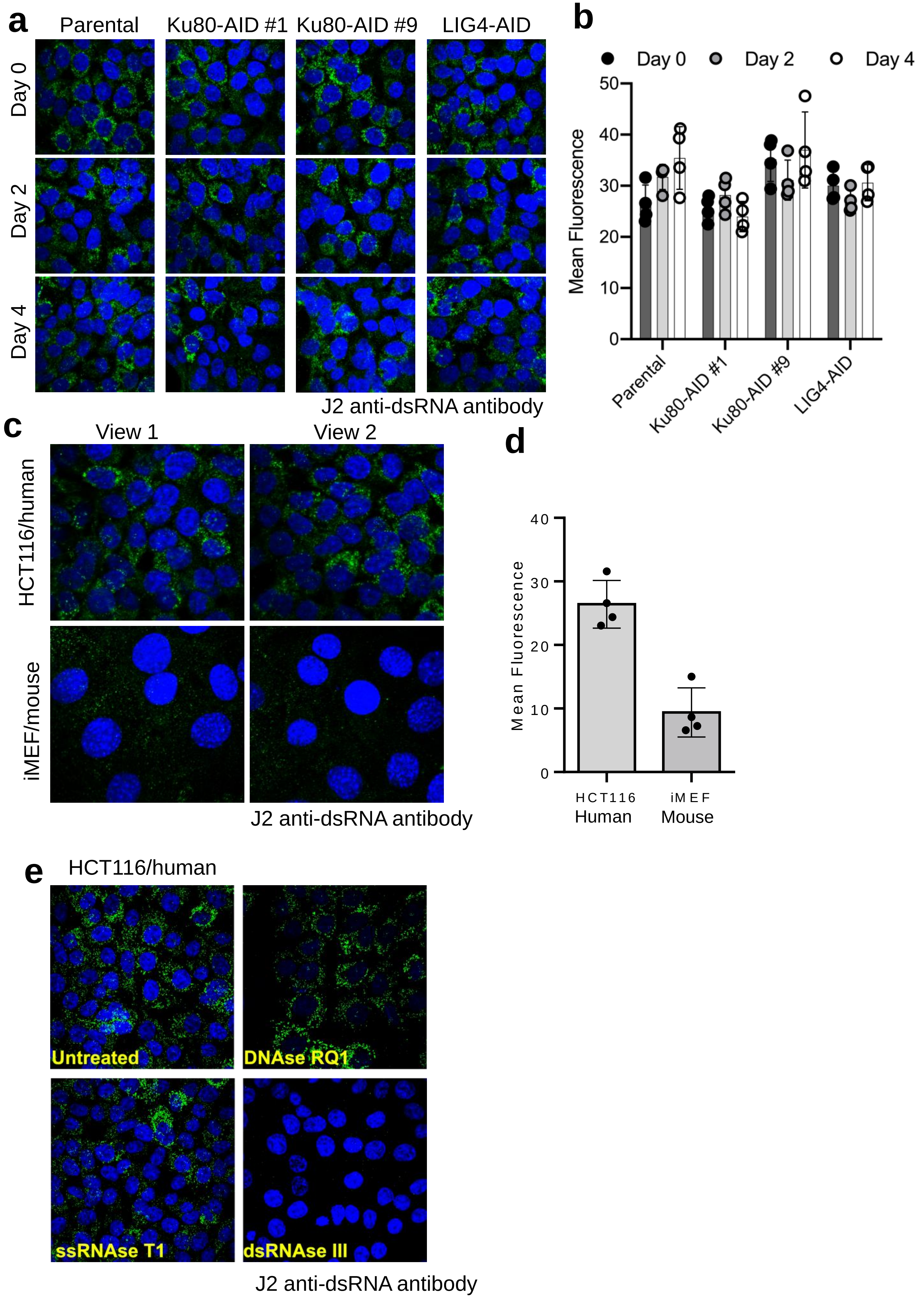

Extended data Fig. 17

**a** tRNA-Lys-AAA (chr17:8,022,473-8,022,548)

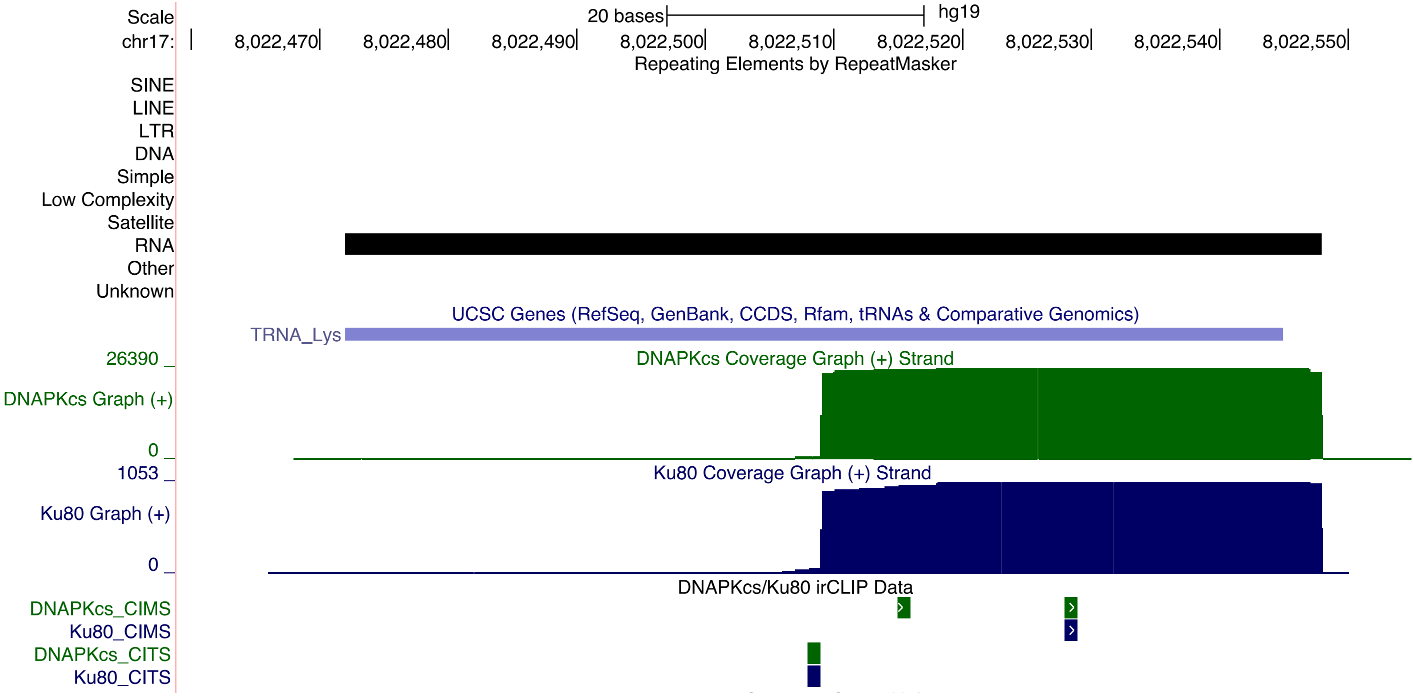


**b** tRNA-Cys-TGY (chr14:73,429,679-73,429,752)

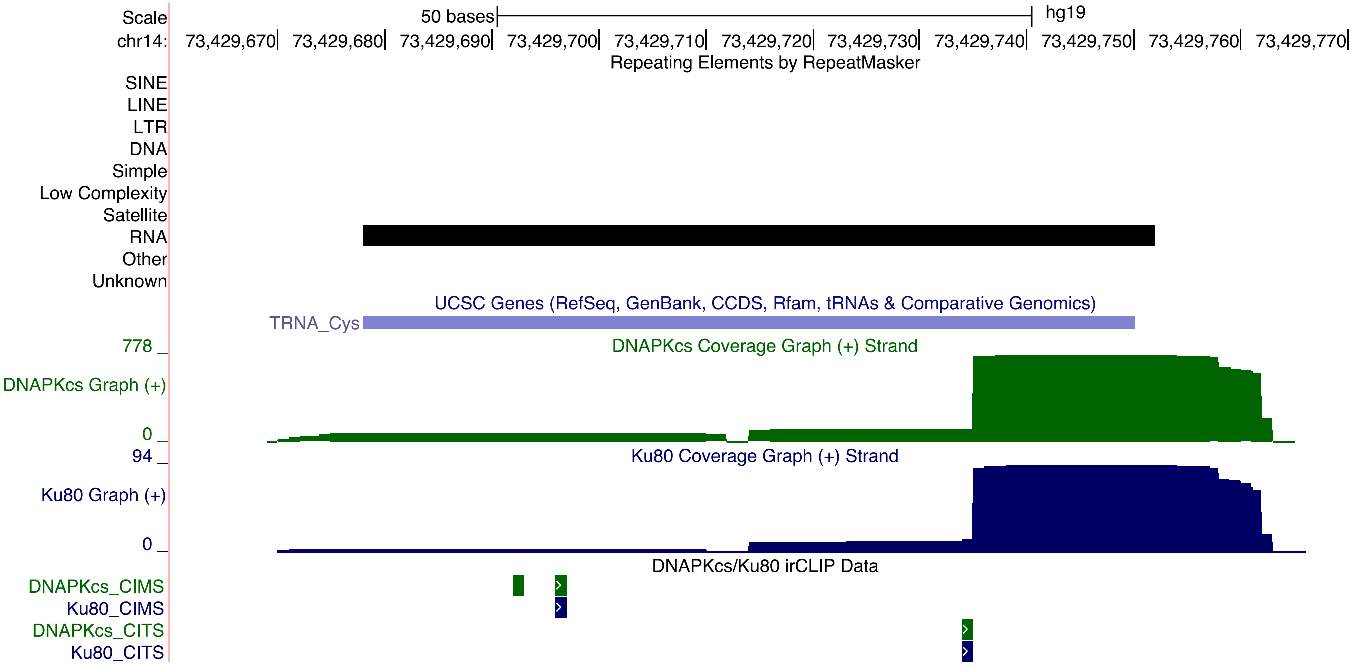

Extended data Fig. 18

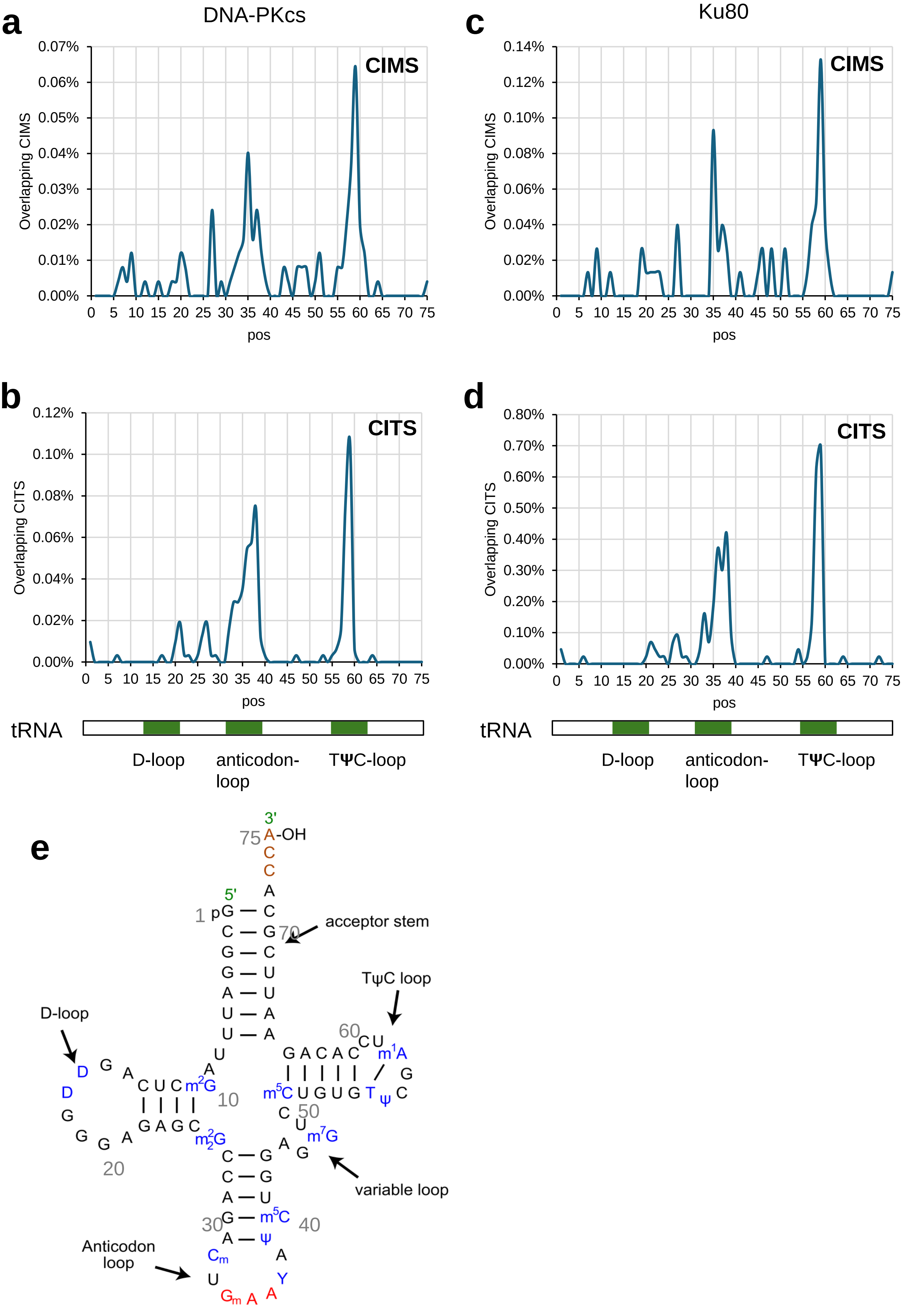
